# Looking Across Tasks: Domain-General and Task-Specific Gaze Behaviours in Spatial Cognition

**DOI:** 10.64898/2026.09.22.753562

**Authors:** Zitong Wu, Daniela E. Aguilar Ramirez, Claudia L. R. Gonzalez

## Abstract

Eye movements have been shown to reveal the cognitive processes underlying spatial ability, but most studies investigating gaze behaviour in spatial cognition use a single, screen-based task. Therefore, it is unknown whether these gaze behaviours are preserved across naturalistic task environments, or whether their relationship with spatial performance is domain-general or task-specific. The present study tested whether established gaze behaviours are preserved and generalizable across diverse, naturalistic spatial tasks. *N* = 30 young adults completed three naturalistic spatial tasks varying in perceptual, cognitive, and visuomotor demands: Q-bitz, the Brick Building Task (BBT), and the Mental Rotation Test (MRT). Gaze was measured using the Pupil Labs Neon eye tracker. *Within-task* analyses were conducted to assess whether gaze behaviours were preserved, and *cross-task* analyses were conducted to assess the generalizability of preserved gaze behaviours. Within-task analyses identified source inspection and transition rate as gaze behaviours that were consistently sensitive to changing task demands. Cross-task analyses showed that source fixation duration was a domain-general predictor of performance, whereas the relationship between transition rate and performance differed by task. Exploratory sex-separated analyses also suggested a more consistent gaze-performance relationship in males. These findings show that source fixation duration may represent a common cognitive process underlying performance across naturalistic spatial tasks.

## 1. Introduction

Spatial ability broadly refers to the capacity to perceive, represent and manipulate spatial information, including the ability to mentally rotate, and transform objects in space (Kolb & Whishaw, 2009). Because it is functionally present in everyday life, spatial ability has broad real-world significance across age groups and contexts. Childhood and adolescent spatial ability strongly predict pursuit of and success in STEM study (Shea et al., 2001; Wai et al., 2009; Gilligan et al., 2017). Spatial ability declines with age and may be an early predictor of cognitive disorders such as Alzheimer’s disease (Iachini et al., 2009; Techentin et al., 2014). Because spatial ability is malleable through training and exposure (Uttal et al., 2013), its measurement is relevant for assessment and skill development in spatially demanding professions such as surgery (Keehner et al., 2004).

Gaze behaviours provide insight into the cognitive processes underpinning spatial performance (Just & Carpenter, 1985; Xue et al., 2017). Gaze metrics assessed in previous eye tracking spatial task studies include pupil diameter, saccade amplitude, fixation count, and fixation duration. Changes in these measures across difficulty conditions and between functionally distinct task regions have been shown to be consistent with different cognitive processes in spatial problem solving, such as cognitive effort, encoding depth, solving strategy, attention allocation, and processing efficiency (Just & Carpenter, 1976, 1985; Khooshabeh et al., 2013; Xue et al., 2017; Nazareth et al., 2019; Toth & Campbell, 2019; Wang et al., 2026). Notably, fixation behaviour directed toward the task source can provide insight into how spatial information is extracted and processed. For example, Just & Carpenter (1980) showed that increased duration of fixations to the task source is linked to deeper processing of the source, which suggests more effortful encoding of the spatial information it contains.

While eye tracking has been utilised extensively for investigation of spatial ability, most studies are conducted on a computerized interface using conventional tasks that isolate spatial cognitive processing (Just & Carpenter, 1976, 1985; Khooshabeh et al., 2013; Xue et al., 2017; Nazareth et al., 2019; Toth & Campbell, 2019; Wang et al., 2026), such as the Mental Rotation Test (Shepard & Metzler, 1971). Additionally, most studies have used only a single task for assessment. This work has been invaluable for the understanding of spatial cognitive strategy, but it remains unclear how well-established gaze behaviours generalize to more naturalistic spatial processing. Such settings can involve physical interaction, visuomotor coordination, and fewer spatial constraints than highly controlled computerized tasks, potentially altering gaze behaviour. Additionally, the single-task design of most studies means that while the cognitive processes associated with different gaze behaviours are relatively well established, it remains unclear whether these behaviours are consistent across different spatial tasks and whether their relationships with spatial performance are preserved. As eye tracking technology advances, its real-world applications will become increasingly relevant. For example, eye tracking has been explored for measurement of cognitive load while driving (Recarte & Nunes, 2003) and during surgical training (Tolvanen et al., 2022). Therefore, it is crucial to understand how gaze behaviours established in computerized spatial tasks translate across spatial tasks with differing real-world demands. Importantly, consistency of a gaze behaviour across tasks does not necessarily mean that its relationship with performance is also preserved. A gaze behaviour may represent a stable individual characteristic across spatial contexts while its functional relevance to performance remains task dependent.

The current study aimed to investigate whether these established gaze behaviours are preserved and generalizable across spatial tasks differing in naturalistic and visuomotor demands. To do this, gaze metrics were assessed across three distinct, spatial tasks: Q-bitz, the Brick Building Task (BBT), and the Mental Rotation Test (MRT). The three tasks differ in perceptual, cognitive and visuomotor demands, providing a continuum of spatial demands across which generalizability can be assessed. Q-bitz involves physically reconstructing a mentally transformed representation within fixed boundaries. Of the three tasks, the BBT is the most naturalistic and demands more free-form planning, construction, and visuomotor coordination with minimal spatial constraint. The MRT utilises pure mental transformation, with motor response limited to circling predetermined answers. Because the MRT has been extensively studied and shown to isolate mental rotation processing, its inclusion serves as an anchor for comparison with the more naturalistic, multi-component tasks. Therefore, gaze behaviours that are consistent across tasks, and show consistent relationships with spatial performance, may reflect domain-general aspects of spatial processing rather than task-specific behaviours.

It was hypothesized that gaze behaviours would remain consistent across tasks, and that at least some of these stable gaze behaviours would show preserved relationships with spatial performance across tasks with differing naturalistic demands. An exploratory component was also included to investigate possible sex differences in generalizability of gaze behaviours. Because a male advantage in mental rotation ability is well established (Voyer et al., 1995; Lauer et al., 2019; Aguilar Ramirez et al., 2021, 2022, 2026), we explored whether gaze behaviours and their cross-task relationships differed between females and males, which may reflect sex-specific strategies for spatial problem solving.

## 2. Methods

### 2.1 Participants

Thirty young adult and late adolescent participants (14 females) took part in the study, ranging in age from 16-27 years old. The sample was restricted to young adults and late adolescents to minimize potential age-related differences in spatial performance (Techentin et al., 2014). All participants were healthy with no diagnosed neurological conditions. Participants who required corrective lenses were eligible to participate if wearing contact lenses, as glasses were incompatible with the Pupil Labs Neon eye tracker.

Participants were recruited through word of mouth within the University of Lethbridge and the broader Lethbridge community, and thus broadly reflect the demographics of the area. Participants identified as White (*n* = 23, 77%), Asian (*n* = 4, 13%), and Mixed Race (*n* = 3, 10%). All recruited participants self-identified as cisgender. Written informed consent was collected from all participants, including participants under the age of 18 who were assessed as having capacity to provide informed consent. The study was approved by the University of Alberta Research Ethics Board (Pro00134357).

### 2.2 Q-bitz

In the Q-bitz task, participants reproduced a target pattern using 16 identical wooden cubes (Aguilar Ramirez et al., 2026). Each participant completed 12 unique trials across three conditions – four trials reproducing a blank control (Figure 1), four trials directly reproducing a target pattern (Figure 2), and four trials in which participants viewed a target pattern and were instructed to mentally rotate it 90° clockwise before reproducing the resulting rotated pattern (Figure 3). The control (C) condition provided a measure of baseline motor performance, while pattern (P), and rotated pattern (R) conditions provided two levels of spatial transformation demand. The P condition of Q-bitz has been shown to strongly relate to MRT performance (Aguilar Ramirez et al., 2026) and the R condition was included to further increase mental rotation load.

**Figure 1:**
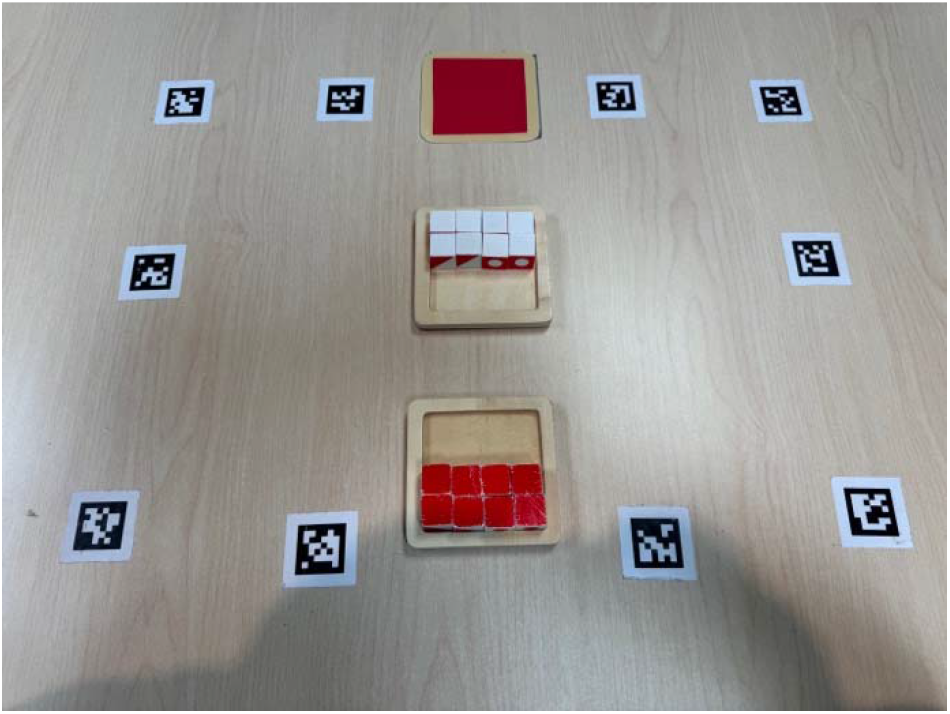
Q-bitz Control (C) condition with lower half of replication completed.

**Figure 2:**
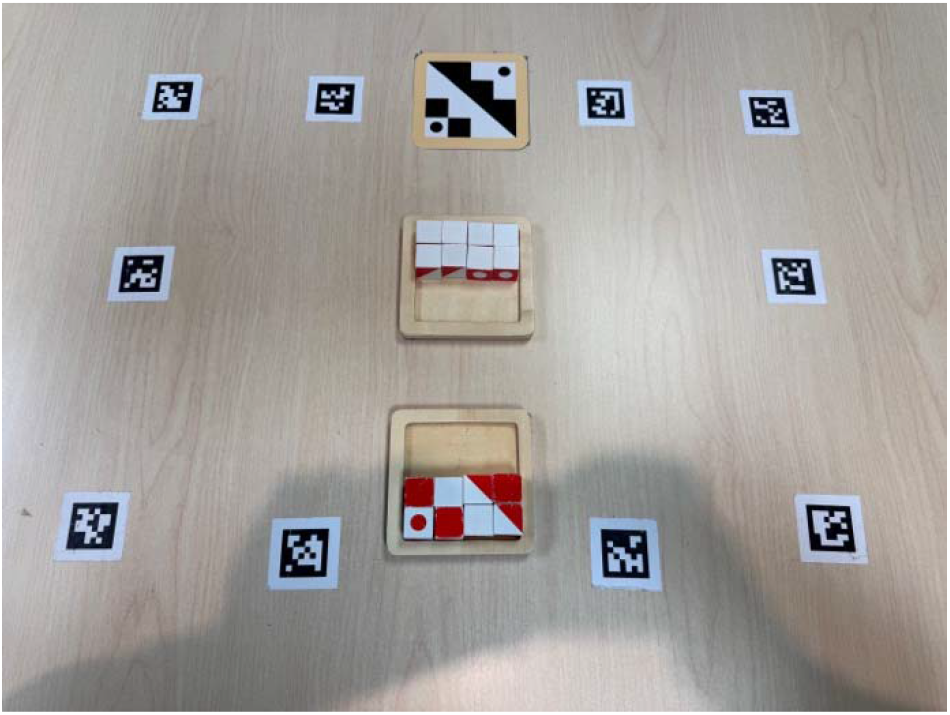
Q-bitz Pattern (P) condition with lower half of replication completed.

**Figure 3:**
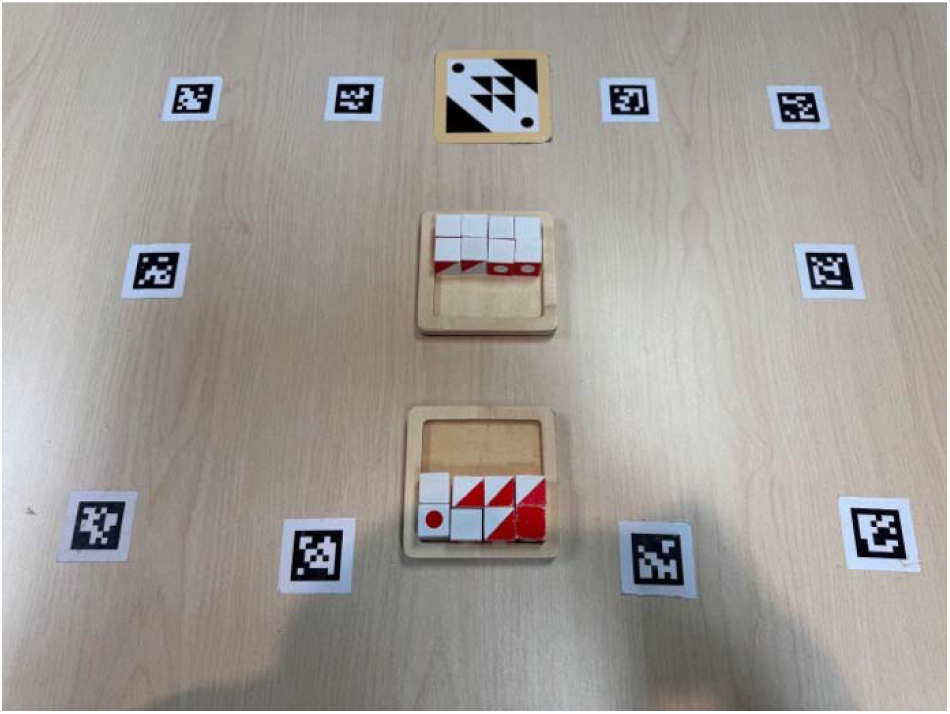
Q-bitz Rotated pattern (R) condition with lower half of replication completed.

Participants were given 30 seconds to complete as much as possible of each trial with only one hand. Within each condition, two trials were completed with the right hand and two with the left hand. The use of one hand increased task difficulty and reduced the likelihood of participants completing the pattern before the end of the 30-sec period.

This standardized task duration and removed the need of time-normalization for eye tracking measures.

### 2.3 Brick Building Task (BBT)

In the Brick Building Task (BBT; Aguilar Ramirez et al., 2020), participants recreated two Lego models: a low mental rotation (LMR; Figure 4) and high mental rotation (HMR; Figure 5) condition, each containing the same 12 bricks. The LMR model was designed to be largely visible in a single plane, while the HMR model rearranged the bricks in a highly three-dimensional arrangement. This provided two levels of mental rotation demand while controlling for the materials used. The BBT has been shown to produce significant performance differences between LMR and HMR conditions, as well as correlations with MRT performance that suggest shared spatial mechanisms (Aguilar Ramirez et al., 2020, 2021, 2022; Pasescu et al., 2026). The participant’s brick supply was standardized as 60 bricks (48 correct pieces, 12 distractors) arranged in an identical layout between participants (Figure 6).

**Figure 4:**
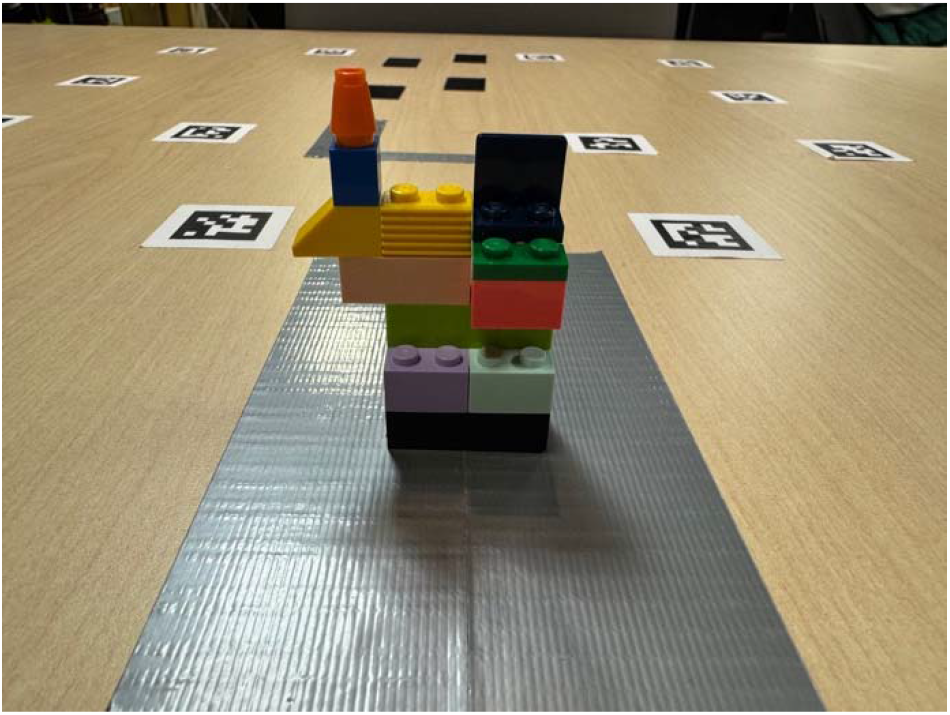
low mental rotation (LMR) model of the Brick Building Task (BBT)

**Figure 5:**
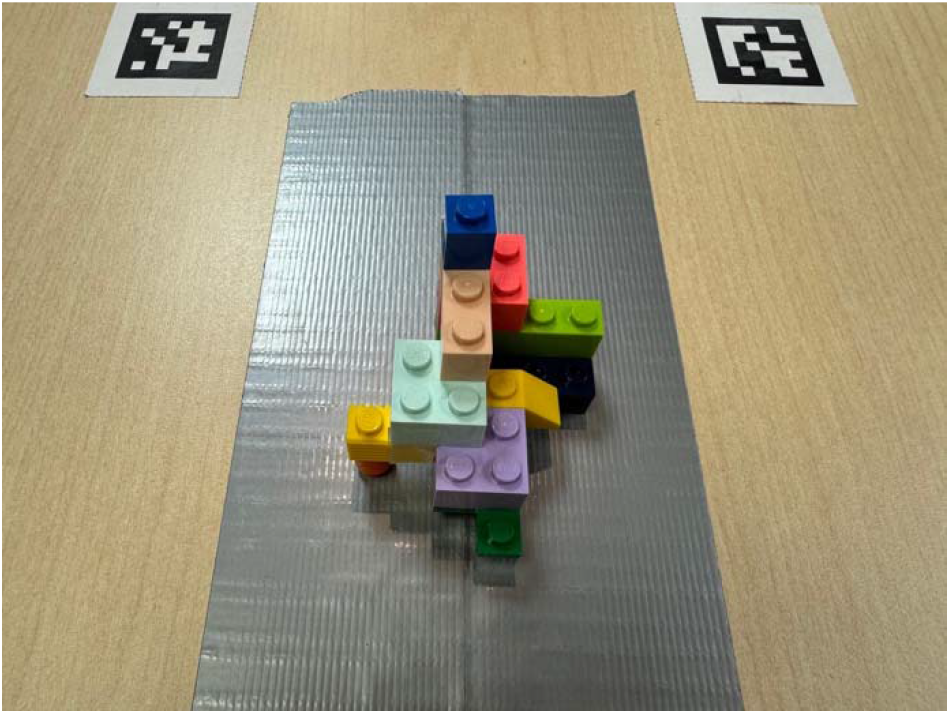
high mental rotation (HMR) model of the Brick Building Task (BBT)

**Figure 6:**
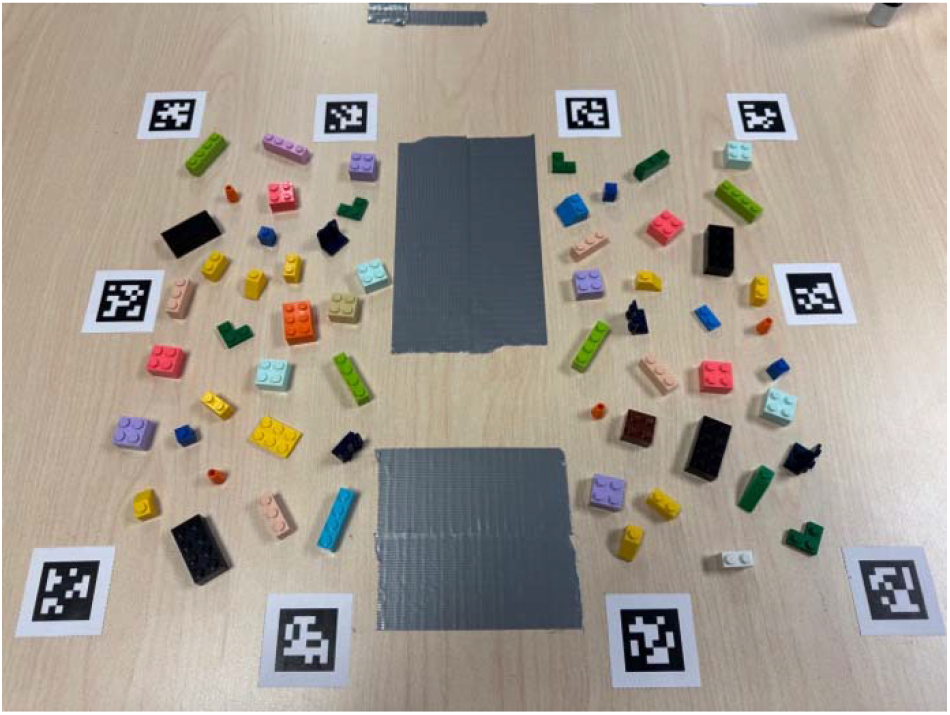
brick supply and layout of the Brick Building Task (BBT)

### 2.4 Mental Rotation Test (MRT)

The Mental Rotation Test (MRT) included the first 12 items from MRT(A) (Peters et al., 1995). Each item contained a reference shape on the left side and four comparison shapes on the right (Figure 7). Participants were required to identify the two comparison figures that represented the correct rotated versions of the reference figure. Participants were given three minutes to complete the set of 12 items.

**Figure 7:**
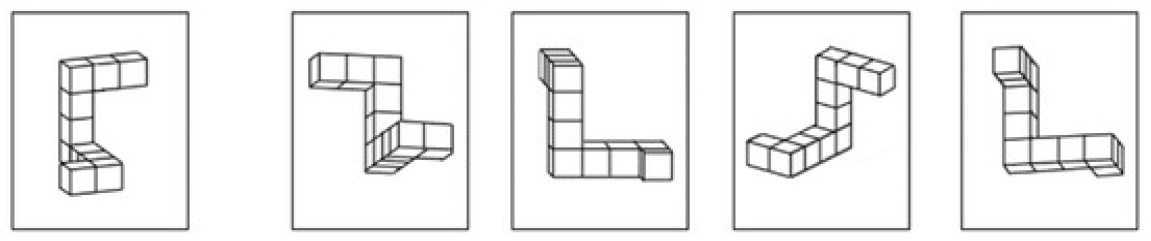
this example illustrates one item from the Mental Rotation Test (MRT). In the example shown, the correct choices are the second and third figures (adapted from Aguilar Ramirez et al., 2026).

To facilitate accurate eye tracking, each MRT item was enlarged to fit a landscape-oriented, letter size sheet of paper. All items were laminated and presented to the participant in a stack (Figure 8)

**Figure 8:**
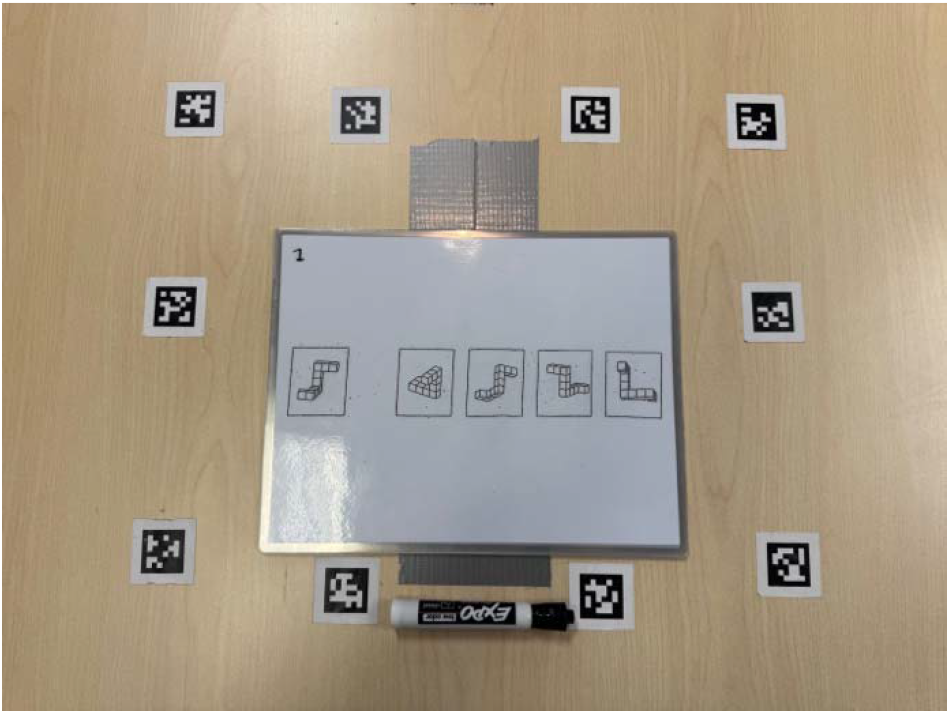
layout of the Mental Rotation Test (MRT)

### 2.5 Procedure

#### 2.5.1 General procedures

Participants were asked to complete an informed consent form and a demographics form before taking part in the study. The Pupil Labs Neon eye tracker glasses were used for eye tracking across all tasks. Ten AprilTags were positioned on the table surrounding the working space, and task layouts were standardized across participants for consistency of eye tracking measures (Figures 1-8). Before beginning the tasks, the Neon eye tracker was adjusted for the participant’s interpupillary distance in the Neon Companion app. Though the Neon eye tracker utilizes a deep learning gaze estimation model that does not require calibration, gaze accuracy was verified using the Offset Correction function in Neon Companion (Pupil Labs, n.d.) by having participants fixate on each of the four corner AprilTags.

Fifteen individual eye tracking videos were recorded in total for each participant – 12 Q-bitz trials, two BBT trials, and a single three-minute video for all MRT trials. At the beginning of each video, participants were first instructed to look around the working space for five seconds before the subject of the task was revealed, at which point they could begin. This period was used to establish a baseline pupil diameter prior to task onset and was designed to minimize differences in factors such as surface luminance and viewing distance, which can influence pupil diameter (Winn et al., 2018). However, previous spatial ability research that successfully isolated cognitive load-mediated pupil dilation largely involved a computer workspace (Campbell et al., 2018; Toth & Campbell, 2019; Wang et al., 2026), and this isolation is more difficult in a naturalistic environment. Therefore, pupillometry measures were treated as exploratory.

The order of task presentation (Q-bitz, BBT, and MRT) was counterbalanced across participants. Within tasks, the easier and harder conditions within Q-bitz and BBT were counterbalanced, but MRT item order remained constant. The order of target pattern cards within Q-bitz was randomly generated for each participant.

#### 2.5.2 Q-bitz procedure

For the Q-bitz task, participants were instructed to replicate the pattern card for each trial as quickly and accurately as possible in 30 seconds, using either their right or left hand. In the rotated pattern condition, they were instructed to mentally rotate the target pattern card 90° clockwise and reproduce the resulting rotated pattern. Prior to each trial, the participant was instructed to look between the cube supply and replication space for five seconds to establish the pre-task pupil baseline (Figures 1-3). The pattern card was then revealed and the 30 second timer was started.

#### 2.5.3 BBT procedure

In the BBT, the participant was instructed to keep the model in the farther grey square and their replication in the closer grey square (Figure 6) to differentiate the two areas for eye tracking. They were advised to replicate the model as quickly and accurately as possible using the bricks provided. The participant was required to turn away during setup of each BBT trial due to the controlled location of each brick in the working space. Before each model was revealed, the participant looked in circles around the building pieces for five seconds to capture pupil baseline measures. Once presented with the model, they were free to move and rotate it within its grey square but could not lift it by more than two inches. Once replicated, the participant’s verbal confirmation was required to denote task completion.

#### 2.5.4 MRT procedure

For the MRT, participants were instructed to identify two of the four figures that matched the reference figure for each item. Participants were given three minutes to complete the set of 12 items and were advised to prioritize accuracy over speed as only items with both answers correct would be counted. Before beginning the task, the figures of the first item were covered with a brightness-matched laminated sheet of paper, and the participant was instructed to look in circles around the white page for the pupil baseline measurement. The three-minute timer was started as the first item was revealed, and the researcher removed each item as it was completed to reveal the next item underneath. The participant was instructed not to move the stack of items as their consistent position allowed for more accurate tracking of gaze behaviour to each area of interest.

### 2.6 Behavioural Data

#### 2.6.1 Q-bitz behavioural data

In each trial, Q-bitz score was calculated as the number of cubes placed correctly with a maximum possible score of 16. The four trials within each condition were then averaged into a single value (*totalcorrect*). Separate analyses were conducted for the individual P and R trials to directly assess differences between the ‘easy’ and ‘hard’ conditions. To assess whether sex differences in baseline motor performance warranted normalizing to the control condition (C) for between-participant analysis, an independent sample T-test was conducted to assess sex differences in total cubes placed in C, which reflects baseline motor speed. No significant difference was found between females (*M* = 13.61, *SD* = 1.71) and males (*M* = 13.02, *SD* = 1.51); (*t*(28) = 1.01, *p* = 0.32). Therefore, there was no evidence of a sex difference in baseline motor speed, and the control condition was not used for normalization. The average score of the rotated pattern condition (R) was also divisively normalized relative to the average score of the regular pattern condition (P), providing a measure of performance in R relative to P (RnP). Because R and P shared the same baseline motor requirements, the control condition was not included in this relative measure.

#### 2.6.2 BBT behavioural data

In the BBT, time to completion of each condition was used to measure performance (*time*). The number of errors made in each condition (*errors*) was also considered as a potential measure of performance but was not analyzed further due to the low number of errors observed.

#### 2.6.3 MRT behavioural data

The MRT was scored as the sum of only the trials in which both answers were correct, with a maximum possible score of 12 (*MRTscore*). An accuracy score was also calculated, which took *MRTscore* as a percentage of the total items completed (*MRT%score*). Both score and accuracy were included to distinguish overall performance from accuracy among attempted items, as participants varied in the number of items completed. The time of each trial was also measured. Since not all participants completed the same number of trials, we could not analyse individual trials to differentiate ‘easy’ and ‘hard’ conditions like the Q-bitz and BBT. Instead, trials were combined to form two outcome groups for each participant and data measure: average of correct trials and average of incorrect trials. Previous work with the MRT has demonstrated that task difficulty, quantified by angular disparity of the rotated figures, scales with both cognitive load and error rate (Bauer et al., 2022). However, because difficulty was not experimentally manipulated in the present MRT, correct and incorrect trials were not treated as direct equivalents of the ‘easy’ and ‘hard’ conditions in Q-bitz and BBT. Comparisons between correct and incorrect trials were therefore treated as exploratory. Within the behavioural data, only trial times could be grouped by averages of correct and incorrect trials (*averagetime*).

### 2.7 Eye Tracking Data

#### 2.7.1 General eye tracking data

Eye tracking videos were processed in Pupil Labs’ Pupil Cloud service. Markers were added to denote the true beginning of each trial, separating it from the pupil baseline measure at the start of each video. Areas of interest (AOIs) were defined to measure gaze behaviour in specific areas of each task (Figures 9-11).

**Figure 9:**
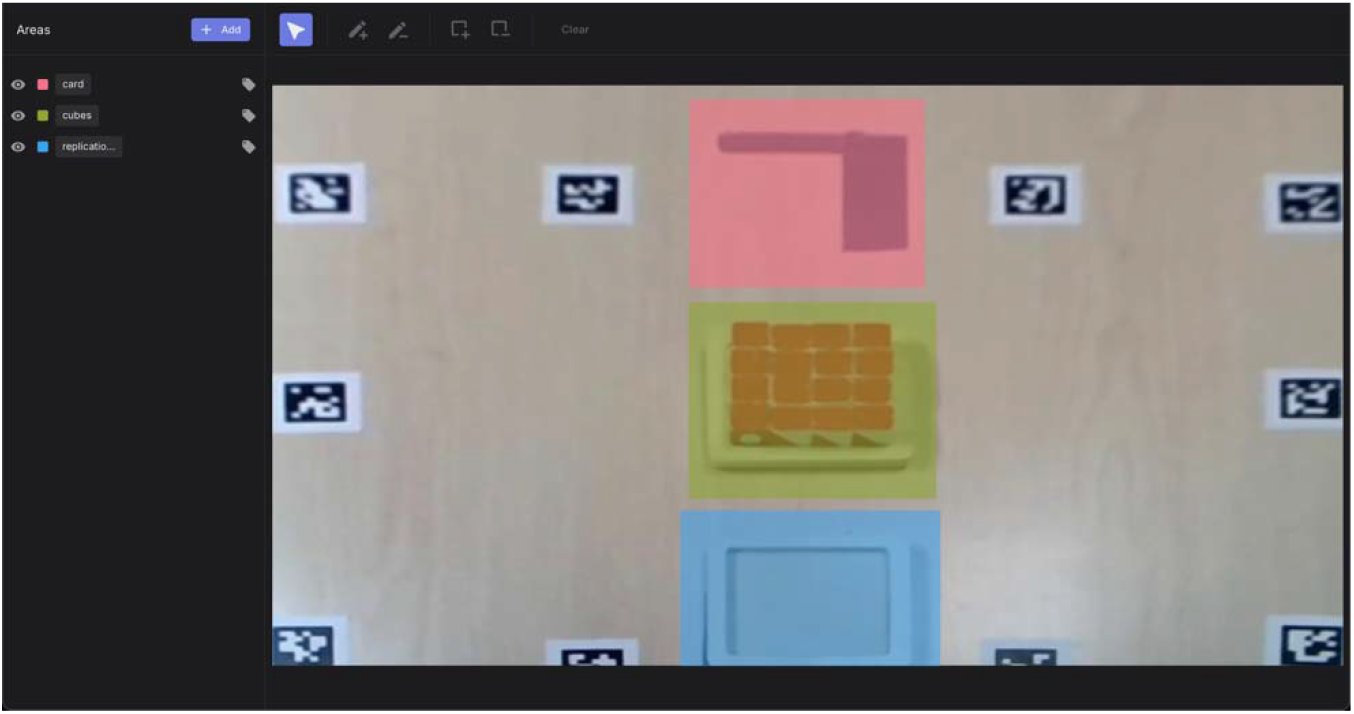
area of interest (AOI) mapping for Q-bitz in Pupil Cloud.

**Figure 10:**
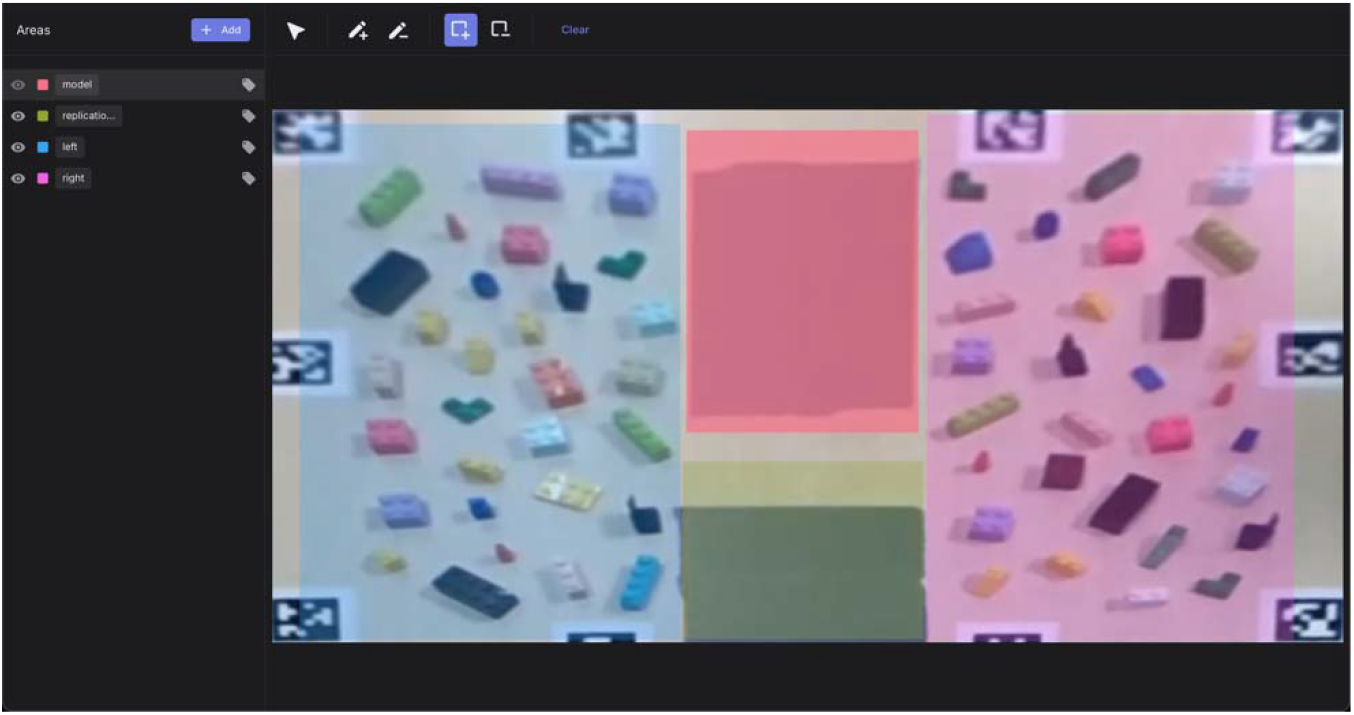
area of interest (AOI) mapping for BBT in Pupil Cloud.

**Figure 11:**
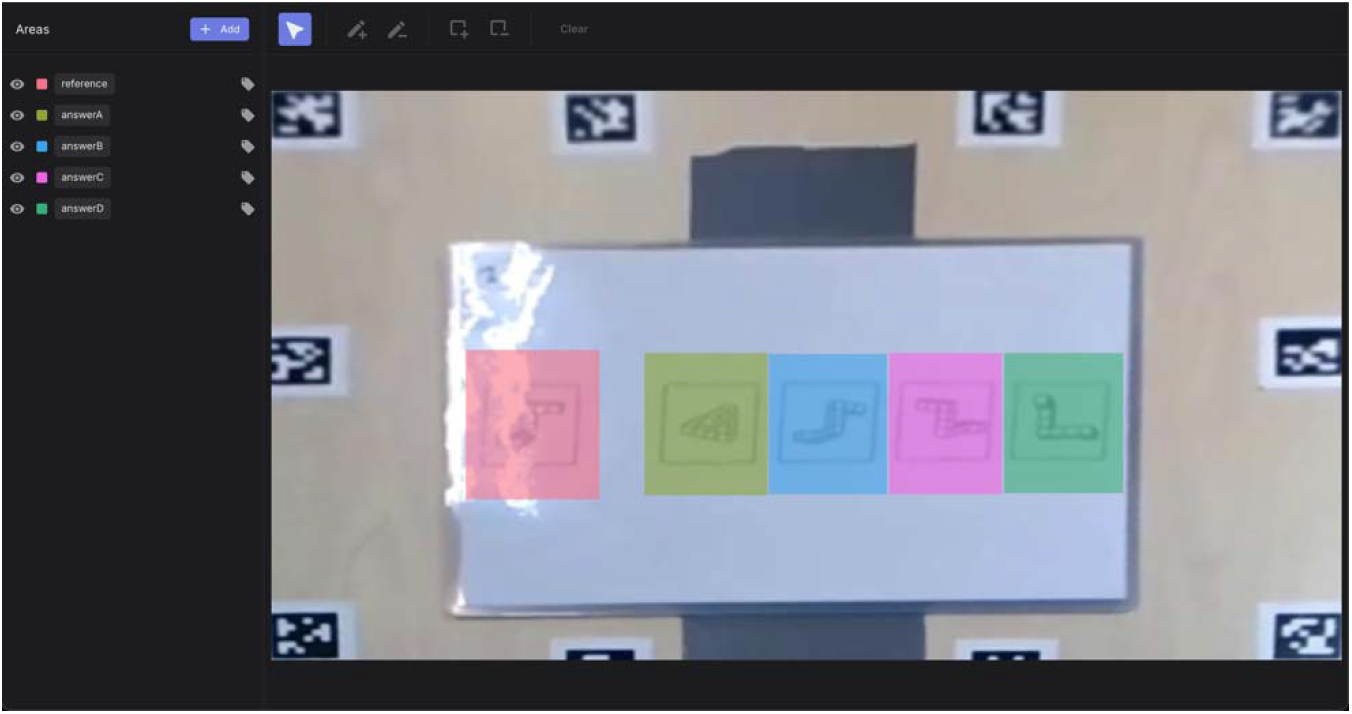
area of interest (AOI) mapping for MRT in Pupil Cloud.

The following eye tracking measures were assessed for all tasks:

– *pupildiameter*: median of pupil diameters from 0.5-2s into the trial, subtractively normalized to the median pupil diameter 1s before the trial. Exploratory.
– *saccadeamplitude*: median saccade amplitude of the whole task in degrees.
– *fixationproportion*: fractional proportion of gaze allocation to each AOI.
– *fixationduration*: median duration (milliseconds) of each individual fixation to an AOI, used as an index of processing duration per glance (Just & Carpenter, 1980).
– *AOItransitions*: transition rate of fixations between AOIs, calculated by the number of fixation transitions from one AOI to another normalized to trial duration.

This provided an index of the rate of switching between task-relevant areas. *fixationproportion*, *fixationduration*, and *AOItransitions* were measured for the unique areas of interest in each task.

#### 2.7.2 Q-bitz eye tracking data

Q-bitz AOIs measured included the pattern card (*card*), the supply of cubes (*cubes*), and the replication space (*replication*) (Figure 9). Transitions were measured between all AOIs (*AOItransitionstotal*) and to and from the pattern card AOI (*AOItransitionscard*). Since Q-bitz trials were restricted to 30 seconds, *AOItransitions* were not time-normalized but remained proportional to transition rate. Like the behavioural data, each eye tracking measure was individually analysed for R and P and also taken as the value of R normalized to P, to quantify the relative change in gaze behaviour between the two conditions. *pupildiameter* and *saccadeamplitude* were normalized divisively to best visualize the proportion of change between conditions. Because all *fixationproportion* measures sum to 1, they were normalized subtractively to preserve this relationship. All fixation measures were subsequently normalized subtractively for consistency.

#### 2.7.3 BBT eye tracking data

BBT AOIs measured included the space containing the model (*model*), the replication space (*replication*), and a combination of the left– and right-side brick supplies (*allpieces*) (Figure 10). Transitions were measured between all AOIs (*AOItransitionstotal*) and to and from the model (*AOItransitionsmodel*). *AOItransitions* measures were excluded from within-task correlation and regression analyses and cross-task mixed-effects models because BBT performance was assessed using trial time and the transition measures were themselves normalized to trial time, creating mathematical dependence between the variables. BBT eye tracking measures remained separate for each condition (LMR, HMR) and all non-transition measures were not normalized.

#### 2.7.4 MRT eye tracking data

MRT AOIs measured included the reference figure (*reference*), the two correct answers for each trial (*truecorrect*), and the two incorrect answers for each trial (*trueincorrect*) (Figure 11). Transitions were measured between all AOIs (*AOItransitionstotal*), to and from the reference AOI (*AOItransitionsreference*), and between the four answer AOIs (*AOItransitionsanswers*). Like the behavioural data, each eye tracking measure was averaged separately across correct and incorrect trials.

Due to the extensive volume of eye tracking data produced, extraction, normalization, and organization of raw eye tracker data was carried out using researcher-verified code developed with the assistance of a large language model (Claude Opus 4.8, high effort reasoning mode). All calculations and data organization were subsequently verified by the researchers.

### 2.8 Data analysis

#### 2.8.1 Within-task data analysis methods

Within each task, a 2×2 mixed-design analysis of variance (ANOVA) was conducted for all dependent variables, with condition (P vs R in Q-bitz, LMR vs HMR in BBT, and correct vs incorrect trial outcomes in MRT) as a within-subjects factor and sex (female vs male) as a between-subjects factor. All significant main effects were supplemented with descriptive statistics, and significant interactions were followed up with simple main effects analyses.

To assess associations between task performance and eye tracking measures, Pearson correlations (*r*) and stepwise multiple linear regressions were conducted for each task. Correlations were calculated first to characterise pairwise associations and are reported in appendices for completeness. Stepwise multiple regressions were then used as exploratory within-task analyses to assess the combined contribution of eye tracking measures to performance. Within each task, both analyses were separated by condition (P vs R in Q-bitz, LMR vs HMR in BBT, correct vs incorrect trial outcome in MRT) in all participants. Each condition-separated analysis was also separated by sex. Due to small sample sizes (*n* = 13-16 per sex), all sex-split analyses were underpowered and susceptible to overfitting in regression. They are therefore included on an exploratory basis to characterise potential sex differences to be explored with future work.

Regressions within all tasks were performed using a stepwise method due to the exploratory nature of the analyses and absence of a priori hypotheses for predictors.

For Q-bitz, in addition to each condition (P and R), correlation analyses were also separately calculated for RnP. To reduce overlap among regression predictors, *fixationproportioncard* was the only *fixationproportion* measure included, and *AOItransitionstotal* was the only *AOItransitions* measure included.

To reduce overlap among regression predictors in the BBT, *fixationproportionmodel* was the only *fixationproportion* measure included. *AOItransitions* measures were only included in the mixed-design ANOVA because they are normalized to trial time and were therefore excluded from analyses predicting trial time to avoid mathematical dependence between the predictor and outcome. Two female and three male participants were excluded from BBT *pupildiameter* analyses due to a task-specific change in normalization protocol during early data collection. Two male participants were excluded from all BBT analyses involving fixation data (*fixationduration*, *fixationproportion*, *AOItransitions*) due to a technical error in AOI mapping that yielded no usable fixation data.

For the MRT, one-way between-subjects ANOVAs were also used to assess sex differences in *MRTscore* and *MRT%score*. To reduce overlap among regression predictors, *fixationproportiontruecorrect* was the only *fixationproportion* measure included, and *AOItransitionstotal* was excluded due to overlap with *AOItransitionsreference* and *AOItransitionsanswers*. Q-bitz and BBT tasks only included

*AOItransitionstotal* because it encompassed more data than the source-specific transition measure (*card*, *model*). MRT is the only task with three *AOItransitions* measures, two of which do not overlap. The two non-overlapping measures were included instead of *AOItransitionstotal* to maximise the detail of the analyses. *averagetime* was also excluded to isolate eye behaviour predictors. Separate sets of regression analyses were conducted to predict *MRTscore* and *MRT%score* and to explore potential sex differences in gaze–performance relationships. Two female and three male participants were excluded from MRT *pupildiameter* analyses due to a task-specific change in normalization protocol during early data collection. Three male participants completed the MRT with no incorrect answers and were thus excluded from correct vs incorrect analyses.

#### 2.8.2 Cross-task data analysis methods

In addition to all within-task analyses, cross-task analyses were conducted to assess generalizability of gaze behaviours. Only measures with functionally equivalent areas of interest between tasks and AOI-independent measures could be considered. Therefore, *fixationproportion* and *fixationduration* were each grouped by source (Q-bitz *card*, BBT *model*, and MRT *reference*) and replication (Q-bitz *replication*, BBT *replication*, and MRT *answers*). The two measures together were further conceptually grouped as source inspection and replication inspection, because both measure visual engagement with the area of interest. In each task, rate of transitions involving the source (Q-bitz *AOItransitionscard*, BBT *AOItransitionsmodel*, and MRT *AOItransitionsanswers*) were also considered together. *pupildiameter*, *saccadeamplitude*, and transition rate (*AOItransitionstotal*) were considered as global, AOI-independent measures. Of these measures, only those which showed consistent associations with difficulty and/or performance across within-task analyses were considered. Measures of fixation duration and proportion to resource (Q-bitz *cubes*, BBT *allpieces*) were not considered for cross-task analyses due to the absence of an equivalent AOI in the MRT.

Cross-task correlation analyses were first performed for task performance to assess whether individual performance remained consistent across tasks. Cross-task correlation analyses were also performed for gaze measures shared across tasks to assess consistency of gaze behaviours across tasks. Linear mixed-effects models were performed to assess whether shared gaze measures predicted performance consistently across tasks. If gaze behaviour remains consistent across tasks, and if this generalized gaze pattern can also consistently predict performance across tasks, this would suggest a common pattern of visual behaviours associated with performance across distinct spatial tasks. All cross-task analyses were also exploratorily separated by sex.

Pearson correlations (*r*) were calculated for all cross-task correlations of task performance (Q-bitz *totalcorrect*, BBT *time*, *MRTscore*). For consistency, each measure was z-scored and BBT time was reverse-scored so higher values denoted stronger performance, directionally matching the other tasks. For each eye tracking measure, Pearson correlations (*r*) were calculated for all combinations of tasks (Q-bitz/BBT, Q-bitz/MRT, BBT/MRT).

In the mixed-effects models, eye tracking measures were averaged across all conditions for each task, yielding a single metric per participant per task for conversion to z-score. Since the number of correct and incorrect MRT trials differed across participants, the MRT eye tracking measures were weighted by the number of correct and incorrect trials. All eye tracking and performance measures (Q-bitz *totalcorrect*, BBT *time*, and *MRTscore)* were standardized to z-scores within the task, with the BBT scored in reverse so higher values denoted stronger performance. This placed all tasks on a common scale for direct comparison. For models examining AOI transitions, BBT was excluded because transition measures were normalized to trial time, which was also the BBT performance measure. Each model included one or more gaze predictors, task, and each predictor’s interaction with task as fixed effects, with task coded using sum contrasts, sex as a covariate, and a by-participant random intercept to account for the non-independence of observations from the same participant. Each predictor’s main effect indexed its pooled association with performance, while its predictor × task interaction tested whether that association differed across tasks. A significant main effect together with a non-significant interaction was interpreted as evidence of an overall association with performance with no detected evidence that the association differed across tasks. Models were estimated using restricted maximum likelihood and fixed effects were evaluated using F-tests with Satterthwaite-approximated degrees of freedom.

Given the study’s exploratory scope and large number of statistical tests, analyses were primarily interpreted as hypothesis-generating and no corrections were applied to the broad correlation, regression, and sex-split analyses. Holm-Bonferroni correction was applied within each effect (across measures) of each within-task ANOVA and within each cross-task mixed-effects model, as these analyses most directly addressed the primary study questions. Effects that did not survive correction are noted and interpreted with caution.

JASP 0.95.0 (Apple Silicon) was used for all statistical analyses.

## 3. Results

### 3.1 Q-bitz within-task analyses

#### 3.1.1 Q-bitz ANOVAs

The results of the mixed-design ANOVAs (difficulty × sex) for Q-bitz are presented in Table 1.

**Table 1:** mixed-design ANOVA results for Q-bitz (difficulty × sex) with significant values (p < .05) bolded.

| Measure | df | Difficulty (P vs R) |  |  | Sex (female vs male) |  |  | Difficulty × Sex |  |  |
| --- | --- | --- | --- | --- | --- | --- | --- | --- | --- | --- |
| | | *F* | *p* | $\eta^2$ | *F* | *p* | $\eta^2$ | *F* | *p* | $\eta^2$ |
| *totalcorrect* | 1, 28 | **212.55** | **< .001\*** | **.88** | 0.03 | .871 | .00 | **6.03** | **.021** | **.18** |
| *pupildiameter* | 1, 28 | 0.04 | .850 | .00 | 2.04 | .164 | .07 | 0.92 | .346 | .03 |
| *saccadeamplitude* | 1, 28 | 1.05 | .314 | .04 | 0.55 | .464 | .02 | 3.03 | .093 | .10 |
| *fixationproportioncard* | 1, 28 | **58.58** | **< .001\*** | **.68** | 0.03 | .859 | .00 | 0.06 | .815 | .00 |
| *fixationproportioncubes* | 1, 28 | **55.28** | **< .001\*** | **.66** | 0.02 | .904 | .00 | 0.18 | .676 | .01 |
| *fixationproportionreplication* | 1, 28 | 3.90 | .058 | .12 | 0.00 | .959 | .00 | 0.33 | .570 | .01 |
| *fixationdurationcard* | 1, 28 | **7.26** | **.012** | **.21** | 2.03 | .165 | .07 | 0.73 | .400 | .03 |
| *fixationdurationcubes* | 1, 28 | **11.72** | **.002\*** | **.30** | **8.80** | **.006** | **.24** | **6.12** | **.020** | **.18** |
| *fixationdurationreplication* | 1, 28 | **16.60** | **< .001\*** | **.37** | **14.89** | **< .001\*** | **.35** | 1.02 | .321 | .04 |
| *AOItransitiontotal* | 1, 28 | **33.49** | **< .001\*** | **.55** | **4.21** | **.050** | **.13** | 0.18 | .671 | .01 |
| *AOItransitionscard* | 1, 28 | 0.99 | .327 | .03 | **5.42** | **.027** | **.16** | 1.36 | .254 | .05 |

Note. Holm-Bonferroni correction was applied within each effect (across measures); asterisks (\*) denote effects that survived correction.

**Table 1:** descriptive statistics for all significant main effects of Q-bitz mixed-design ANOVA.

| Significant Measure | Pattern (P) |  | Rotated (R) |  | Female |  | Male |  |
| --- | --- | --- | --- | --- | --- | --- | --- | --- |
| | $\mu$ | *SD* | $\mu$ | *SD* | $\mu$ | *SD* | $\mu$ | *SD* |
| *totalcorrect* | 10.37 | 1.98 | 6.91 | 2.22 |  |  |  |  |
| *fixationproportioncard* | 0.346 | 0.040 | 0.434 | 0.062 |  |  |  |  |
| *fixationproportioncubes* | 0.322 | 0.054 | 0.257 | 0.046 |  |  |  |  |
| *fixationproportionreplication* | 0.332 | 0.051 | 0.309 | 0.053 |  |  |  |  |
| *fixationdurationcard* | 320.90 | 75.39 | 355.96 | 95.30 |  |  |  |  |
| *fixationdurationcubes* | 278.23 | 39.40 | 257.83 | 43.34 | 249.00 | 38.43 | 284.68 | 27.15 |
| *fixationdurationreplication* | 331.25 | 64.28 | 284.84 | 57.78 | 275.29 | 39.62 | 336.70 | 46.57 |
| *AOItransitiontotal* | 43.77 | 7.31 | 38.84 | 8.57 | 44.20 | 8.12 | 38.77 | 6.38 |
| *AOItransitionscard* |  |  |  |  | 34.02 | 7.15 | 29.00 | 4.52 |

Descriptive statistics are presented in Table 2 to clarify the direction of each significant main effect.

**Table 2:** significant results of regression analyses for Q-bitz Pattern (P) and Rotated pattern (R) condition measures.

| Dependent variable/condition | Retained predictor(s) | B | SE B | $\beta$ | t | p |
| --- | --- | --- | --- | --- | --- | --- |
| P – All – *totalcorrect* | *AOItransitiontotal* | 0.176 | 0.039 | .65 | 4.56 | < .001 |
| P – Females – *totalcorrect* | *AOItransitiontotal* | 0.157 | 0.048 | .63 | 3.24 | .008 |
|  | *pupildiameter* | 5.969 | 2.623 | .44 | 2.28 | .044 |
| P – Males – *totalcorrect* | *AOItransitiontotal* | 0.254 | 0.064 | .73 | 3.96 | .001 |
| R – All – *totalcorrect* | *AOItransitiontotal* | 0.162 | 0.029 | .63 | 5.53 | < .001 |
|  | *fixationdurationcubes* | 0.029 | 0.006 | .57 | 5.29 | < .001 |
|  | *fixationdurationcard* | -0.007 | 0.003 | -.32 | -2.97 | .006 |
|  | *fixationproportioncard* | -10.844 | 3.703 | -.30 | -2.93 | .007 |
| R – Females – *totalcorrect* | *AOItransitiontotal* | 0.174 | 0.041 | .70 | 4.28 | .001 |
|  | *fixationdurationcubes* | 0.027 | 0.009 | .48 | 2.96 | .013 |
| R – Males – *totalcorrect* | *AOItransitiontotal* | 0.226 | 0.051 | .76 | 4.43 | < .001 |

Simple main effect analyses were calculated for significant interactions. For *totalcorrect*, the simple main effect of difficulty was significant in both females (*F*(1, 13) = 165.77, *p* < .001, η² = .93) and males (*F*(1, 15) = 68.14, *p* < .001, η² = .82). This was due to a larger P-to-R decline in females (Δ = –4.08) than males (Δ = –2.91), indicating a greater rotation cost in females. For *fixationdurationcubes*, the simple main effect of difficulty was significant in females (*F*(1, 13) = 16.44, *p* = .001, η² = .56) but not males (*F*(1, 15)= 0.48, *p* = .499, η² = .03). Descriptive statistics show that on average, female fixation durations to cubes shortened with difficulty (Δ = –37.0), while the change was smaller in males (Δ = –6.0) and fixation durations were higher in males overall (Table 2).

#### 3.1.2 Q-bitz correlations and regressions

Pearson correlation coefficients (*r*) of all measures were calculated for Pattern (P) condition with all participants (Figure A1), females only (Figure A2), and males only (Figure A3), as well as for Rotated pattern (R) condition with all participants (Figure A4), females only (Figure A5), and males only (Figure A6). Separate correlations were calculated for the R normalized to P condition of Q-bitz with all participants (Figure A7), females only (Figure A8), and males only (Figure A9).

Stepwise regression analyses were performed to assess retained predictors of performance in each correlation category, separately for P and R (Table 3) and R normalized to P (Table 4).

**Table 3:** significant results of regression analyses for Q-bitz R normalized to P (RnP) condition measures.

| Dependent variable/condition | Retained predictor(s) | *B* | *SE B* | $\beta$ | *t* | *p* |
| --- | --- | --- | --- | --- | --- | --- |
| RnP – All – *totalcorrect* | *AOItransitiontotal* | 0.016 | 0.004 | .51 | 4.11 | < .001 |
| | *fixationdurationcubes* | 0.001 | $4.902 \times 10^{-4}$ | .28 | 2.24 | .034 |
|  | *fixationproportioncard* | -0.897 | 0.295 | -.38 | -3.04 | .005 |
| | *fixationdurationcard* | $-5.413 \times 10^{-4}$ | $2.577 \times 10^{-4}$ | -.27 | -2.10 | .046 |
| RnP – Females – *totalcorrect* | *AOItransitiontotal* | 0.017 | 0.004 | .80 | 4.61 | < .001 |
| RnP – Males – *totalcorrect* | *AOItransitiontotal* | 0.028 | 0.006 | .57 | 4.33 | .001 |
| | *fixationdurationcubes* | 0.001 | $5.705 \times 10^{-4}$ | .29 | 2.21 | .049 |
|  | *fixationproportioncard* | -1.040 | 0.268 | -.49 | -3.87 | .003 |
| | *fixationdurationcard* | $-7.385 \times 10^{-4}$ | $2.160 \times 10^{-4}$ | -.43 | -3.42 | .006 |

**Table 4:** mixed-design ANOVA results for the BBT (difficulty × sex) with significant values (p < .05) bolded. *Note*. Holm-Bonferroni correction was applied within each effect (across measures); asterisks (*) denote effects that survived correction.

| Measure | *df* | Difficulty (LMR vs HMR) | | | Sex (female vs male) | | | Difficulty × Sex | | |
| --- | --- | --- | --- | --- | --- | --- | --- | --- | --- | --- |
| | | *F* | *p* | $\eta^2$ | *F* | *p* | $\eta^2$ | *F* | *p* | $\eta^2$ |
| *time* | 1, 28 | **99.18** | **< .001\*** | **.78** | 0.04 | .851 | .00 | 0.17 | .681 | .01 |
| *errors* | 1, 28 | 1.88 | .181 | .06 | 2.18 | .151 | .07 | 0.01 | .928 | .00 |
| *pupildiameter* | 1, 23 | 0.08 | .775 | .00 | **11.93** | **.002\*** | **.34** | 2.63 | .119 | .10 |
| *saccadeamplitude* | 1, 28 | **12.71** | **.001\*** | **.31** | 0.01 | .940 | .00 | 1.63 | .212 | .06 |
| *fixationproportionmodel* | 1, 26 | **30.83** | **< .001\*** | **.54** | 1.38 | .251 | .05 | 0.00 | .995 | .00 |
| *fixationproportionreplication* | 1, 26 | **47.54** | **< .001\*** | **.65** | **7.00** | **.014** | **.21** | 0.01 | .932 | .00 |
| *fixationproportionallpieces* | 1, 26 | **98.11** | **< .001\*** | **.79** | 1.59 | .219 | .06 | 0.01 | .938 | .00 |
| *fixationdurationmodel* | 1, 26 | **10.80** | **.003\*** | **.29** | 3.11 | .090 | .11 | 0.39 | .537 | .02 |
| *fixationdurationreplication* | 1, 26 | 0.51 | .484 | .02 | 2.21 | .149 | .08 | **10.16** | **.004\*** | **.28** |
| *fixationdurationallpieces* | 1, 26 | 0.30 | .586 | .01 | **4.41** | **.046** | **.15** | 3.25 | .083 | .11 |
| *AOItransitiontotal* | 1, 26 | **53.27** | **< .001\*** | **.67** | 1.26 | .272 | .05 | **4.40** | **.046** | **.15** |
| *AOItransitionmodel* | 1, 26 | **23.36** | **< .001\*** | **.47** | 0.66 | .424 | .03 | 2.62 | .118 | .09 |

The P condition model with all participants was significant (*F*(1, 28) = 20.82, *p* < .001, *R*^2^ = .43, adj. *R*^2^ = .41) and positively predicted by *AOItransitionstotal*. With participants separated by sex, the female P condition model was significant (*F*(2, 11) = 7.75, *p* = .008, *R*^2^ = .59, adj. *R*^2^ = .51) and positively predicted by *AOItransitionstotal* and *pupildiameter*. The male P condition model was significant (*F*(1, 14) = 15.67, *p* = .001, *R*^2^ = .53, adj. *R*^2^ = .49) and positively predicted by *AOItransitionstotal*.

The R condition model with all participants was significant (*F*(4, 25) = 20.12, *p* < .001, *R*^2^ = .76, adj. *R*^2^ = .73) and positively predicted by *AOItransitionstotal* and *fixationdurationcubes*, and negatively predicted by *fixationdurationcard* and *fixationproportioncard*. The female R condition model was significant (*F*(2, 11) = 13.48, *p* = .001, *R*^2^ = .71, adj. *R*^2^ = .66) and positively predicted by *AOItransitionstotal* and *fixationdurationcubes*. The male R condition model was significant (*F*(1, 14) = 19.58, *p* < .001, *R*^2^ = .58, adj. *R*^2^ = .55) and positively predicted by *AOItransitionstotal*.

Overall, Q-bitz performance across both conditions (P and R) was consistently predicted by fixation transition count (*AOItransitionstotal*); participants with more transitions scored higher. In the R condition only, gaze behaviour to the card and cube supply also significantly predicted performance. Higher *totalcorrect* was predicted by increased fixation duration to the cubes, and decreased fixation duration and total proportion to the card. In other words, better performers in the R condition fixated less on the target pattern card overall and their fixations were shorter.

The RnP models with all participants (*F*(4, 25) = 10.93, *p* < .001, *R*^2^ = .64, adj. *R*^2^ = .58), females only (*F*(1, 12) = 21.23, *p* < .001, *R*^2^ = .64, adj. *R*^2^ = .61), and males only (*F*(4, 11) = 15.80, *p* < .001, *R*^2^ = .85, adj. *R*^2^ = .80) were all significant. Each retained predictor is in the same direction as when present in the un-normalized R regressions.

In the Q-bitz RnP condition, all predictors aligned with the un-normalized R condition. This suggests that these gaze behaviours were associated with the increased spatial transformation demands of the R condition relative to P. In the male participant regressions, these R-associated predictors are present in the RnP condition but not the un-normalized R condition.

### 3.2 BBT within-task analyses

#### 3.2.1 BBT ANOVAs

The results of the mixed-design ANOVAs (difficulty × sex) for the BBT are presented in Table 5.

**Table 5:**
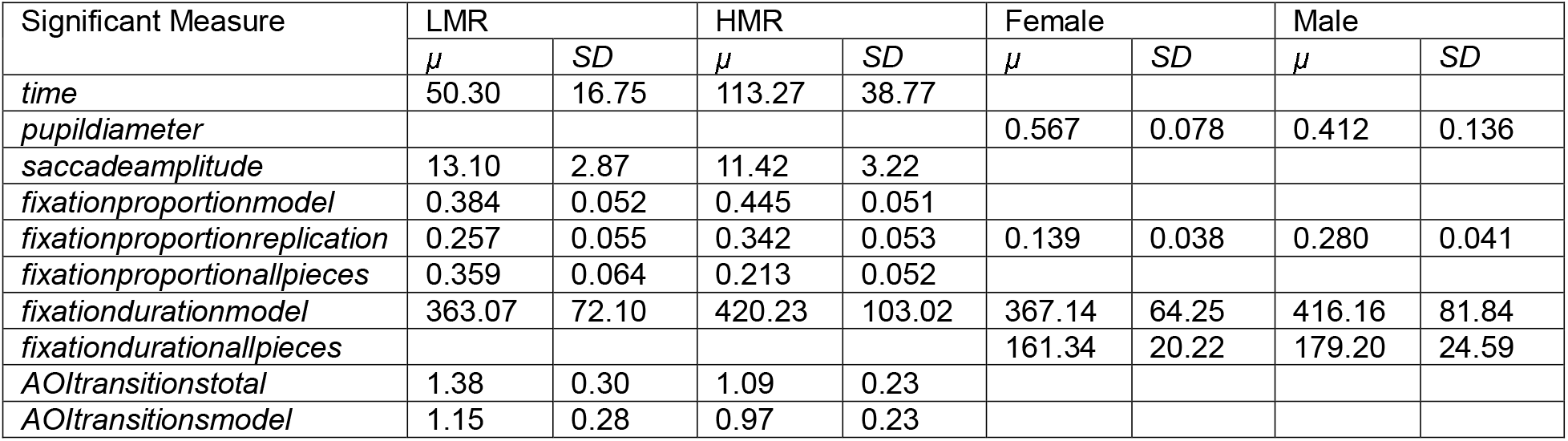
descriptive statistics for all significant main effects of BBT mixed-design ANOVA.

Descriptive statistics are presented in Table 6 to clarify the direction of each significant main effect.

**Table 6:** significant results of regression analyses for BBT LMR and HMR condition measures.

| Dependent variable/condition | Retained predictor(s) | *B* | *SE B* | $\beta$ | *t* | *p* |
| --- | --- | --- | --- | --- | --- | --- |
| LMR – All – *time* | *saccadeamplitude* | -2.739 | 1.007 | -.49 | -2.72 | .012 |
| LMR – Females – *time* | *saccadeamplitude* | -5.058 | 1.813 | -.66 | -2.79 | .019 |
| LMR – Males – *time* | *pupildiameter* | 32.58 | 13.339 | .59 | 2.44 | .033 |
| HMR – All – *time* | None |  |  |  |  |  |
| HMR – Females – *time* | None |  |  |  |  |  |
| HMR – Males – *time* | None |  |  |  |  |  |

Simple main effect analyses were calculated for significant interactions. For *fixationdurationreplication*, the simple main effect of difficulty was significant in females (*F*(1, 13) = 10.16, *p* = .007, η² = .44) but not in males (*F*(1, 13) = 2.45, *p* = .142, η² = .16). Fixation duration to the replication increased from LMR to HMR in females (Δ = 47.2), whereas males showed a numerical decrease (Δ = –30.0). For *AOItransitionstotal*, the simple main effect of difficulty was significant in both females (*F*(1,13) = 37.19, *p* < .001, η² = .74) and males (*F*(1,13) = 16.64, *p* = .001, η² = .56). Descriptive statistics reveal that transition rate decreased from LMR to HMR in both sexes, with a larger decrease in females (Δ = –0.37) than males (Δ = –0.20).

#### 3.2.2 BBT correlations and regressions

Pearson correlation coefficients (*r*) of all measures were calculated for LMR condition with all participants (Figure B1), females only (Figure B2), and males only (Figure B3), as well as for HMR condition with all participants (Figure B4), females only (Figure B5), and males only (Figure B6).

Stepwise regression analyses were performed to assess retained predictors of performance in each correlation category, separately for LMR and HMR (Table 7).

The LMR condition model with all participants was significant (*F*(1, 23) = 7.40, *p* = .012, *R*^2^ = .24, adj. *R*^2^ = .21), with time negatively predicted by *saccadeamplitude*. With participants separated by sex, the female LMR condition model was significant (*F*(1, 10) = 7.79, *p* = .019, *R*^2^ = .44, adj. *R*^2^ = .38), with time negatively predicted by *saccadeamplitude*. The male LMR condition model was significant (*F*(1, 11) = 5.97, *p* = .033, *R*^2^ = .35, adj. *R*^2^ = .29), with time positively predicted by *pupildiameter*. No eye-tracking measures were retained as predictors of time in the HMR condition models.

### 3.3 MRT within-task analyses

#### 3.3.1 MRT ANOVAs

The results of the mixed-design ANOVAs (trial outcome × sex) for each MRT measure are presented in Table 8.

**Table 8:**
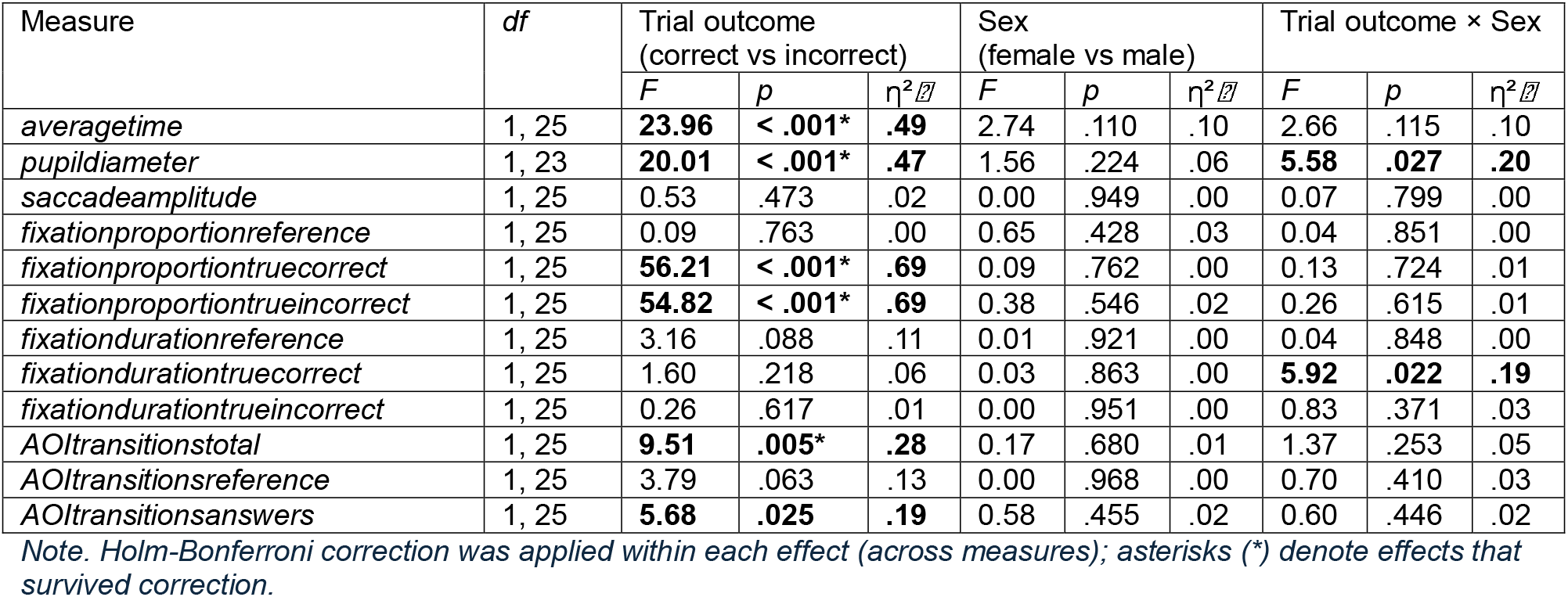
mixed-design ANOVA results for MRT (trial outcome × sex) with significant values (p < .05) bolded.

Descriptive statistics are presented in Table 9 to clarify the direction of each significant main effect.

**Table 9:**
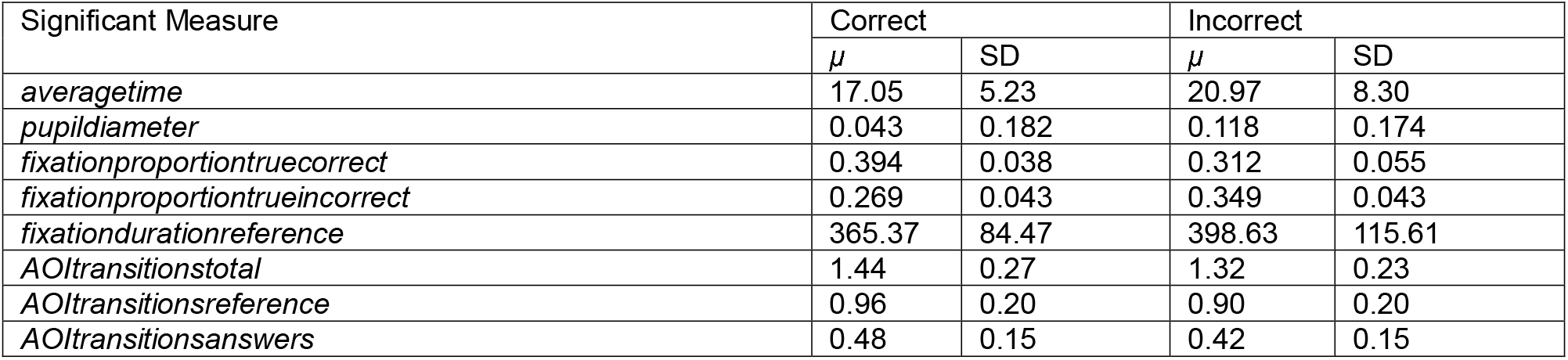
descriptive statistics for all significant main effects of MRT mixed-design ANOVA.

Simple main effect analyses were calculated for significant interactions. For *pupildiameter*, the correct vs incorrect difference was significant in females (*F*(1, 12) = 24.79, *p* < .001, η² = .67) but not in males (*F*(1, 11) = 2.10, *p* = .175, η² = .16). Pupil diameter increased from correct to incorrect trials in both sexes, with a larger increase in females (Δ = 0.112) than males (Δ = 0.034). For *fixationdurationtruecorrect*, the correct vs incorrect difference was significant in females (*F*(1, 13) = 6.91, *p* = .021, η² = .35) but not in males (*F*(1, 12) = 0.68, *p* = .425, η² = .05). Females showed longer fixation durations to the true correct answers during incorrect than correct trials (Δ = 70.7), whereas males showed a small numerical decrease (Δ = –22.4).

Two MRT one-way ANOVAs were conducted with sex as the between-subjects factor, with *MRTscore* and *MRT%score* as the respective dependent variables. No significant sex effects were found with *MRTscore* (*F*(1, 28) = 2.02, *p* = .166, η² = .07), but *MRT%score* revealed a significant main effect of sex (*F*(1, 28) = 11.73, *p* = .002., η² = .30) with males scoring more accurately (higher *MRT%score*). The main effect of sex in *MRT%score* survives Holm-Bonferroni correction (*p_adj* = .004).

#### 3.3.2 MRT correlations and regressions

Pearson correlation coefficients (*r*) of MRTscore, MRT%score, and all measures were calculated for correct trials with all participants (Figure C1), females only (Figure C2), and males only (Figure C3), as well as for incorrect trials with all participants (Figure C4), females only (Figure C5), and males only (Figure C6).

Stepwise regression analyses were performed to assess retained predictors in each correlation category, separately for *MRTscore* (Table 10) and *MRT%score* (Table 11).

**Table 10:**
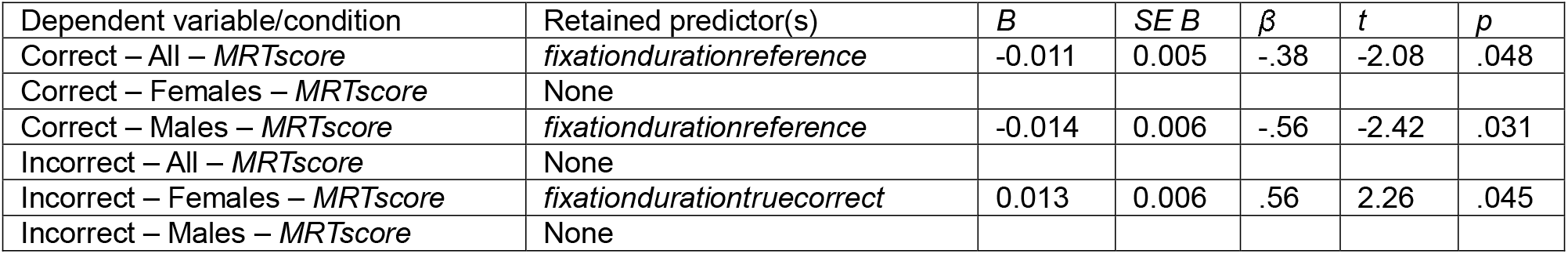
significant results of regression analyses for MRTscore.

**Table 11:** significant results of regression analyses for MRT%score.

| Dependent variable/condition | Retained predictor(s) | *B* | *SE B* | $\beta$ | *t* | *p* |
| --- | --- | --- | --- | --- | --- | --- |
| Correct – All – *MRT%score* | None |  |  |  |  |  |
| Correct – Females – *MRT%score* | None |  |  |  |  |  |
| Correct – Males – *MRT%score* | None |  |  |  |  |  |
| Incorrect – All – *MRT%score* | *AOItransitionsanswers* | -0.551 | 0.223 | -.46 | -2.47 | .021 |
| Incorrect – Females – *MRT%score* | *AOItransitionsanswers* | -1.032 | 0.285 | -.94 | -3.62 | .005 |
|  | *fixationproportiontruecorrect* | 1.901 | 0.842 | .59 | 2.26 | .048 |
| Incorrect – Males – *MRT%score* | *fixationdurationtrueincorrect* | -0.001 | $3.95 \times 10^{-4}$ | -.72 | -3.29 | .008 |

For *MRTscore* with all participants, the correct trials model was significant (*F*(1, 26) = 4.32, *p* = .048, *R*^2^ = .14, adj. *R*^2^ = .11) and negatively predicted by *fixationdurationreference*. With participants separated by sex, the male correct trials model was significant (*F*(1, 13) = 5.85, *p* = .031, *R*^2^ = .31, adj. *R*^2^ = .26) and negatively predicted by *fixationdurationreference*. The female incorrect trials model was significant (*F*(1, 11) = 5.10, *p* = .045, *R*^2^ = .32, adj. *R*^2^ = .26) and positively predicted by *fixationdurationtruecorrect*.

For *MRT%score* with all participants, the incorrect trials model was significant (*F*(1, 23) = 6.10, *p* = .021, *R*^2^ = .21, adj. *R*^2^ = .18) and negatively predicted by *AOItransitionsanswers*. With participants separated by sex, the female incorrect trials model was significant (*F*(2, 10) = 6.56, *p* = .015, *R*^2^ = .57, adj. *R*^2^ = .48) and predicted negatively by *AOItransitionsanswers* and positively by *fixationproportiontruecorrect*. The male incorrect trials model was significant (*F*(1, 10) = 10.84, *p* = .008, *R*^2^ = .52, adj. *R*^2^ = .47) and negatively predicted by *fixationdurationtrueincorrect*.

Overall, higher *MRTscore* was associated with shorter fixation durations to the reference during correct trials. Higher *MRT%score*, which measures accuracy, was associated with a lower rate of fixation transitions between the answers during incorrect trials. Exploratory sex-separated models additionally retained gaze measures to the *truecorrect* and *trueincorrect* answer AOIs as predictors of performance.

### 3.4 Cross-task Analyses

#### 3.4.1 Cross-task performance correlations

Pearson correlation analyses were conducted on the measures of performance between each task (Q-bitz *totalcorrect*, BBT *time*, *MRTscore*) to assess participants’ consistency of performance between tasks in all participants and separated by sex (Table 12).

**Table 12:** cross-task correlations of performance with significant values (p < .05) bolded.

| Participants | Tasks | *N* | *r* | *p* |
| --- | --- | --- | --- | --- |
| All | ***Q-bitz/BBT*** | **30** | **.428** | **.018** |
|  | ***Q-bitz/MRT*** | **30** | **.692** | **< .001** |
|  | ***BBT/MRT*** | **30** | **.466** | **.009** |
| Females | *Q-bitz/BBT* | 14 | .332 | .246 |
|  | ***Q-bitz/MRT*** | **14** | **.581** | **.029** |
|  | *BBT/MRT* | 14 | .422 | .133 |
| Males | ***Q-bitz/BBT*** | **16** | **.509** | **.044** |
|  | ***Q-bitz/MRT*** | **16** | **.818** | **< .001** |
|  | ***BBT/MRT*** | **16** | **.506** | **.045** |

Correlation analyses revealed that across all participants, better performance in each single task was associated with better performance in all tasks. When split by sex, significant performance correlations were observed across all task pairs in males, whereas only the Q-bitz/MRT correlation reached significance in females.

#### 3.4.2 Cross-task gaze behaviour correlations

In previous within-task analyses, transition rate positively predicted performance across all Q-bitz conditions (Tables 3 & 4) and decreased with difficulty in all tasks (Tables 1, 5, & 8). Decreased source inspection also predicted performance across multiple tasks, with Q-bitz R performance predicted by decrease in both fixation duration and proportion to the card (Table 3), and MRT performance predicted by lower fixation duration to the reference in correct trials (Table 10). Fixation duration to the source also increased with difficulty in all tasks (Tables 1, 5, & 8). Therefore, transition rate and source inspection were included in cross-task analyses.

Since these predictors of performance arise across multiple conditions and tasks, Pearson correlation analyses were conducted to assess whether transition rate (*AOItransitionstotal*) or source inspection (*fixationduration* and *fixationproportion* to Q-bitz *card*, BBT *model*, and MRT *reference*) were consistent across tasks (Table 13). Sex-split analyses were also conducted to exploratorily assess sex differences in consistency of gaze behaviour (Table 14).

**Table 13:** cross-task correlations of gaze behaviour in all participants with significant values (p < .05) bolded.

| Gaze Behaviour | Tasks | N | r | p |
| --- | --- | --- | --- | --- |
| Transition rate | **Q-bitz/BBT** | **28** | **.686** | **< .001** |
|  | **Q-bitz/MRT** | **30** | **.508** | **.004** |
|  | **BBT/MRT** | **28** | **.687** | **< .001** |
| Fixation duration to source | **Q-bitz/BBT** | **28** | **.437** | **.020** |
|  | **Q-bitz/MRT** | **30** | **.595** | **< .001** |
|  | **BBT/MRT** | **28** | **.559** | **.002** |
| Fixation proportion to source | Q-bitz/BBT | 28 | .279 | .151 |
|  | Q-bitz/MRT | 30 | -.262 | .162 |
|  | BBT/MRT | 28 | -.327 | .089 |

**Table 14:**
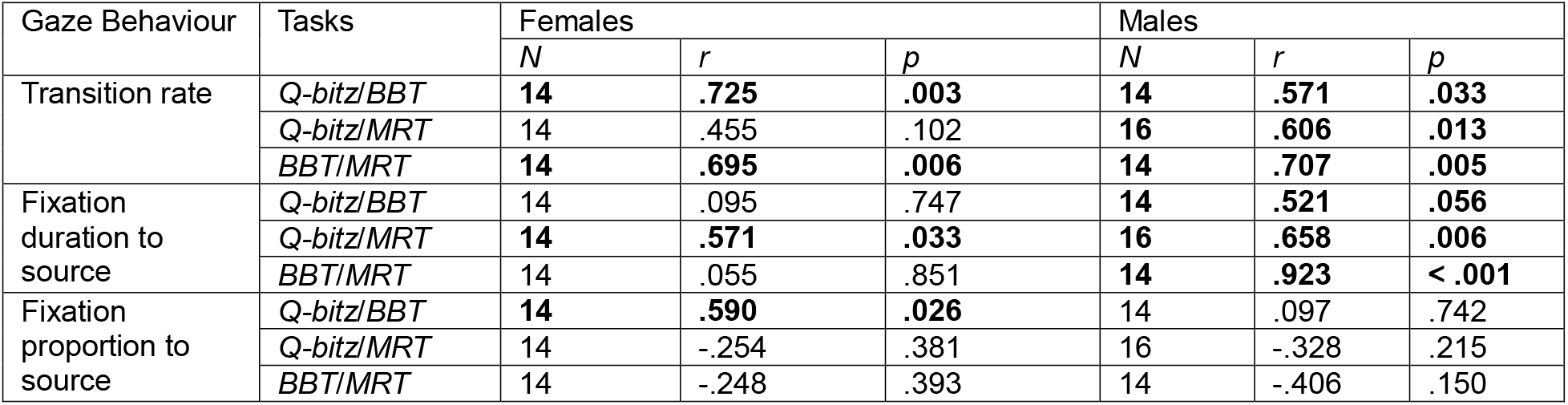
sex-separated cross-task correlations of gaze behaviour with significant values (p < .05) bolded.

| Gaze Behaviour | Tasks | Females |  |  | Males |  |  |
| --- | --- | --- | --- | --- | --- | --- | --- |
|  |  | N | r | p | N | r | p |
| Transition rate | **Q-bitz/BBT** | **14** | **.725** | **.003** | **14** | **.571** | **.033** |
|  | **Q-bitz/MRT** | 14 | .455 | .102 | **16** | **.606** | **.013** |
|  | **BBT/MRT** | **14** | **.695** | **.006** | **14** | **.707** | **.005** |
| Fixation duration to source | **Q-bitz/BBT** | 14 | .095 | .747 | **14** | **.521** | **.056** |
|  | **Q-bitz/MRT** | **14** | **.571** | **.033** | **16** | **.658** | **.006** |
|  | **BBT/MRT** | 14 | .055 | .851 | **14** | **.923** | **< .001** |
| Fixation proportion to source | **Q-bitz/BBT** | **14** | **.590** | **.026** | 14 | .097 | .742 |
|  | **Q-bitz/MRT** | 14 | -.254 | .381 | 16 | -.328 | .215 |
|  | **BBT/MRT** | 14 | -.248 | .393 | 14 | -.406 | .150 |

In all participants, correlation analyses revealed significant relationships between all tasks for transition rate and source fixation duration, indicating that individual differences in these gaze measures were consistent across tasks. When separated by sex, significant correlations across all task pairs were observed in males for transition rate, and source fixation duration was significantly correlated for Q-bitz/MRT and BBT/MRT. Females showed significant transition-rate correlations for Q-bitz/BBT and BBT/MRT, but only the Q-bitz/MRT correlation was significant for source fixation duration.

#### 3.4.3 Cross-task mixed-effects models

Linear mixed-effects models were also used to assess whether these gaze behaviours predicted performance across tasks. Separate models were generated for transition rate (Table 15) and source inspection (Table 16). BBT was removed from the transition rate model because *AOItransitions* and BBT performance were both based on trial time, creating mathematical dependence between variables. For source inspection, both predictors were combined in one model and entered simultaneously to control for each other.

**Table 15:** results of mixed-effects models for transition rate with all participants; with significant values (p < .05) bolded.

| Effect – Transition rate ($N = 30$ participants; 60 observations) | *F* | *df* | *p* | *b* (*SE*) |
| --- | --- | --- | --- | --- |
| ***AOItransitions*** | **12.05** | **1, 51.7** | **.001** | **0.37 (0.11)** |
| ***AOItransitions* × task** | **6.73** | **1, 29.6** | **.015** | **-0.19 (0.08)** |
| Task | 0.00 | 1, 26.2 | n.s. | – |
| Sex | 3.01 | 1, 27.4 | .094 | -0.25 (0.15) |

Note. Random-intercept (participant) variance = 0.48 ( $SD = 0.69$ ); residual variance = 0.26 ( $SD = 0.51$ ). Predictors and outcome standardized within task; sum contrasts, Type III SS, Satterthwaite *df*. All significant effects survived Holm–Bonferroni correction within the model: *AOItransitions* ( $p_{adj} = .004$ ) and *AOItransitions* × task ( $p_{adj} = .045$ ).

**Table 16:**
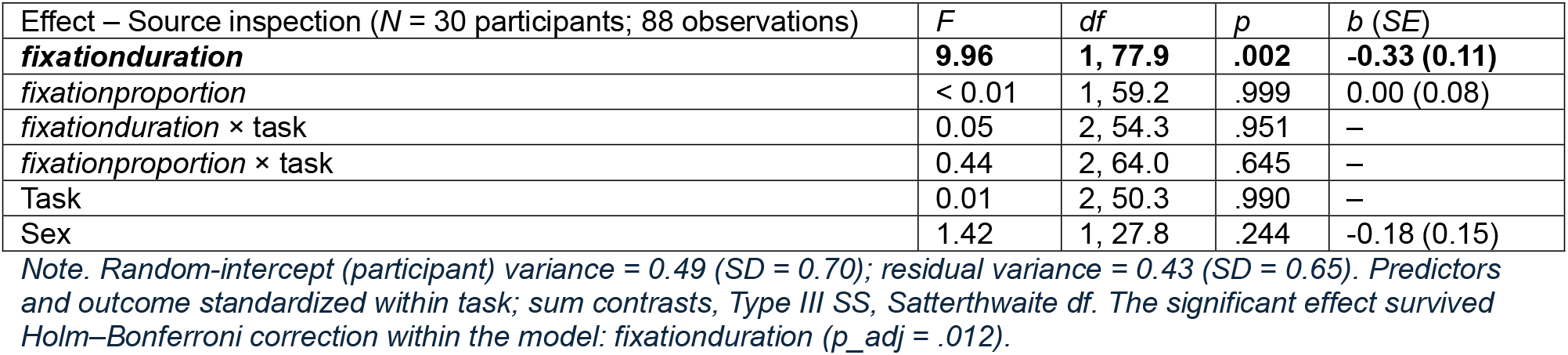
results of mixed-effects models for source inspection with all participants; with significant values (p < .05) bolded.

| Effect – Source inspection ($N = 30$ participants; 88 observations) | *F* | *df* | *p* | *b* (*SE*) |
| --- | --- | --- | --- | --- |
| ***fixationduration*** | **9.96** | **1, 77.9** | **.002** | **-0.33 (0.11)** |
| *fixationproportion* | < 0.01 | 1, 59.2 | .999 | 0.00 (0.08) |
| *fixationduration* × task | 0.05 | 2, 54.3 | .951 | – |
| *fixationproportion* × task | 0.44 | 2, 64.0 | .645 | – |
| Task | 0.01 | 2, 50.3 | .990 | – |
| Sex | 1.42 | 1, 27.8 | .244 | -0.18 (0.15) |

Note. Random-intercept (participant) variance = 0.49 ( $SD = 0.70$ ); residual variance = 0.43 ( $SD = 0.65$ ). Predictors and outcome standardized within task; sum contrasts, Type III SS, Satterthwaite *df*. The significant effect survived Holm–Bonferroni correction within the model: *fixationduration* ( $p_{adj} = .012$ ).

Mixed-effects models with all participants revealed a significant positive main effect of transition rate, indicating that higher transition rates were associated with better performance. However, there was also a significant *AOItransitions* × task interaction, indicating a difference in magnitude of the relationship across tasks. Specifically, simple slope analysis showed that while the relationship was positive in both tasks, *AOItransitions* was a significant predictor of performance in Q-bitz (β = 0.562, *p* < .001) but not in MRT (β = 0.175, *p* = .183). Therefore, the relationship between transition rate and performance differed across tasks despite being directionally consistent. In the source inspection model, *fixationduration* showed a significant negative main effect with no *fixationduration* × task interaction, while *fixationproportion* showed no association. Therefore, shorter source fixation duration was associated with stronger performance across tasks, with no evidence that this relationship differed by task.

Both models were also exploratorily separated by sex (Tables 17 & 18).

**Table 17:** sex-separated results of mixed-effects models for transition rate with significant values (p < .05) bolded.

| Effect – Transition rate | Females (N = 14 participants; 28 observations) |  |  |  | Males (N = 16 participants; 32 observations) |  |  |  |
| --- | --- | --- | --- | --- | --- | --- | --- | --- |
|  | *F* | *df* | *p* | *b* (SE) | *F* | *df* | *p* | *b* (SE) |
| *AOItransitions* | 1.96 | 1, 23.2 | .175 | 0.23 (0.16) | **10.74** | **1, 26.0** | **.003** | **0.49 (0.15)** |
| *AOItransitions* × task | **5.02** | **1, 13.7** | **.042** | **-0.28 (0.13)** | 1.77 | 1, 13.8 | .205 | -0.13 (0.09) |
| Task | 0.02 | 1, 11.8 | .887 | -0.02 (0.12) | 0.01 | 1, 13.4 | .910 | 0.01 (0.08) |

Note. Female random-intercept (participant) variance = 0.46 (SD = 0.67); residual variance = 0.36 (SD = 0.60). Male random-intercept (participant) variance = 0.48 (SD = 0.69); residual variance = 0.18 (SD = 0.43). Predictors and outcome standardized within task; sum contrasts, Type III SS, Satterthwaite *df*.

**Table 18:** sex-separated results of mixed-effects models for source inspection with significant values (p < .05) bolded.

| Effect – Source inspection | Females (N = 14 participants; 42 observations) |  |  |  | Males (N = 16 participants; 46 observations) |  |  |  |
| --- | --- | --- | --- | --- | --- | --- | --- | --- |
|  | *F* | *df* | *p* | *b* (SE) | *F* | *df* | *p* | *b* (SE) |
| *fixationduration* | 2.36 | 1, 31.0 | .135 | -0.28 (0.18) | **5.61** | **1, 35.5** | **.023** | **-0.34 (0.14)** |
| *fixationproportion* | 1.28 | 1, 29.3 | .268 | -0.17 (0.15) | 0.24 | 1, 24.1 | .632 | 0.06 (0.12) |
| *fixationduration* × task | 0.04 | 2, 25.6 | .965 | – | 0.38 | 2, 23.8 | .691 | – |
| *fixationproportion* × task | 0.73 | 2, 27.9 | .492 | – | 1.21 | 2, 29.7 | .312 | – |
| Task | 0.16 | 2, 21.9 | .853 | – | 0.11 | 2, 22.1 | .896 | – |

Note. Female random-intercept (participant) variance = 0.46 (SD = 0.68); residual variance = 0.52 (SD = 0.72). Male random-intercept (participant) variance = 0.50 (SD = 0.71); residual variance = 0.39 (SD = 0.63). Predictors and outcome standardized within task; sum contrasts, Type III SS, Satterthwaite *df*.

Exploratory sex-split mixed-effects models show an absence of all main effects in females, and a significant main effect in both transition rate and source fixation duration in males. Despite this, females showed a significant *AOItransitions* × task interaction because *AOItransitions* was a significant positive predictor of performance in Q-bitz (β = 0.512, *p* = .018) but not in MRT (β = –0.054, *p* = .804). This interaction mirrors the interaction seen with all participants included. Conversely, males showed a main effect of *AOItransitions* but no interaction. These sex-separated patterns are descriptive and should not be interpreted as evidence of a sex difference without a direct statistical test of sex moderation, particularly given the small sample sizes.

## 4. Discussion

### 4.1 Preserved and domain-general gaze properties of spatial cognition

#### 4.1.1 Summary

The present study utilized multiple naturalistic spatial tasks with differing perceptual, cognitive, and visuomotor demands to determine whether established gaze behaviours are task– and environment-specific, or preserved and generalizable across spatial contexts. While many of the gaze behaviours in this study have been described in previous eye tracking spatial ability research, most of these studies have been administered in highly controlled, screen-based settings (Just & Carpenter, 1976, 1985; Khooshabeh et al., 2013; Xue et al., 2017; Nazareth et al., 2019; Toth & Campbell, 2019; Wang et al., 2026). Though ecologically valid studies have been conducted, they have typically examined a single spatial task in isolation, limiting inference about whether gaze behaviours extend across different spatial tasks. Determining whether gaze behaviours persist across these differing task demands can help distinguish task-specific visual strategies from more domain-general properties of spatial cognition.

#### 4.1.2 Source inspection

Across all tasks, source inspection (fixation duration to Q-bitz *card*, BBT *model*, and MRT *reference*) repeatedly changed with increased task demands. Specifically, analyses of variance showed longer fixation durations to the source in the more difficult condition of BBT, with weaker evidence for this pattern in Q-bitz after correction for multiple comparisons, and a similar numerical trend in MRT incorrect versus correct trials. Because increased fixation duration has been associated with deeper encoding (Just & Carpenter, 1980), this suggests that greater spatial demands may require more sustained encoding of source information, resulting in longer source fixations across tasks with very different visuomotor demands.

Regression analyses showed that source inspection (both fixation duration and proportion to source) also consistently predicted task performance. Within-task, within-difficulty regressions demonstrated that better performance in the Q-bitz R condition is predicted by decrease in both fixation durations and proportion to the source. This finding was strengthened by its persistence in the Q-bitz R-normalized-to-P (RnP) analysis, suggesting that the relationship is specifically associated with the increased spatial transformation demands of R relative to P. In the MRT, regression analyses showed that performance was also predicted by decreased fixation durations to the source in correct trials. These findings indicate that within each task, lower source inspection predicts better performance. Paas & Van Merriënboer (1993) established a framework that combined task difficulty and cognitive effort into a single measure of efficiency. Viewed within this framework, better performers and participants in the easier condition may require less sustained inspection because they encode the source information needed for spatial transformation more efficiently. Thus, shorter source inspection may reflect a more efficient relationship between cognitive effort and spatial performance.

The within-task analyses demonstrated that source inspection behaviours established in controlled, single-task settings are preserved in more naturalistic spatial tasks. However, this does not establish the extent to which the behaviour is task-specific and modulated by ecological factors. Cross-task analyses were therefore conducted to assess whether source inspection generalizes across spatial contexts. Cross-task correlations of both performance and source fixation duration were significant between all tasks, showing that individual differences in performance and source fixation duration were preserved across tasks. Importantly, the mixed-effects model showed that this shared gaze pattern significantly predicted performance across all tasks, and its effect was robust to multiple-comparison correction. Together, these findings identify source fixation duration as a domain-general predictor of spatial performance. This raises the possibility that efficient encoding of source information represents a common cognitive process underlying performance across otherwise different spatial tasks.

#### 4.1.3 Resource inspection

Within-task trends in resource inspection (fixation duration and proportion to Q-bitz *cubes* and BBT *allpieces*) were also observed. Analyses of variance showed lower fixation proportion to resources in the difficult condition of both Q-bitz (R) and BBT (HMR), as well as lower fixation duration in Q-bitz R. These findings were robust to multiple-comparison correction. Thus, resource inspection decreased as task demands increased. In Q-bitz, regression analyses also showed that higher fixation duration to *cubes* predicts better performance in the R condition. In both analyses, resource inspection behaviour is inverse to source inspection. Resources are likely the least cognitively consequential component of each task because they do not require active mental rotation like the source or replication. Therefore, the decrease in resource inspection may reflect a reallocation of attention toward the source as difficulty increases. Because resource inspection may therefore be partly dependent on source inspection, and because the MRT does not have any equivalent ‘resource’ measure, cross-task associations of resource inspection were not explored. Thus, resource inspection appears to be preserved across the two more naturalistic construction tasks, but whether it generalizes across spatial tasks remains unknown.

#### 4.1.4 Transition rate

Consistent changes in transition rate (*AOItransitionstotal*) were observed with modulation of difficulty across all tasks. Analyses of variance showed lower transition rate in the more difficult condition of Q-bitz (R) and BBT (HMR), with a similar pattern in MRT incorrect compared with correct trials. These findings were robust to multiple-comparison correction. This likely reflects that during easier spatial processing, participants can minimize working memory demand by frequently re-sampling different task areas rather than retaining them internally (Ballard et al., 1995). As difficulty increased, mental transformation load may have surpassed working memory as the primary burden, leading to deeper encoding and more sustained gaze as shown by the increase in source inspection. Together, these findings suggest that transition rate is sensitive to changing cognitive demands across spatial tasks with very different visuomotor requirements.

In regression, all conditions of Q-bitz (P, R, and RnP) showed increased transition rate as a predictor of better performance. However, this predictive relationship was not present in other tasks. This consistent relationship across Q-bitz conditions supports the relevance of transition rate to performance in a naturalistic spatial task (in accordance with the Paas & Van Merriënboer (1993) framework). However, its absence from the regressions of other tasks may suggest that the relationship between transition rate and performance is task-specific, or more sensitive to differing task demands than source inspection.

Cross-task analyses were used to statistically examine the generalizability of transition rate behaviour. Cross-task correlations of transition rate were significant across all three task pairs (Table 13), showing that participants maintained a similar transition rate across tasks. The mixed-effects model, which excluded the BBT to avoid circularity with its time-based performance measure, showed an overall relationship between transition rate and performance across Q-bitz and MRT. However, a transition rate × task interaction showed that *AOItransitionstotal* predicted performance significantly in Q-bitz but not in the MRT. Both the main effect and interaction were robust to multiple-comparison correction. Therefore, transition rate appears to be a stable characteristic of an individual’s gaze behaviour across spatial tasks, but its relationship with performance is task dependent. This raises the possibility that transition rate reflects a general visual sampling tendency whose functional importance depends on the specific cognitive and visuomotor demands of the task.

### 4.2 Exploratory sex divergence in domain-generalizability of gaze properties

Exploratory sex-separated cross-task analyses showed that cross-task generalizability of source inspection and transition rate was more consistently observed in males than in females. In the mixed-effects models, male performance was significantly associated with both source fixation duration and transition rate but did not have a significant transition rate × task interaction like in all participants. Female mixed-effects models had no significant main effects but retained the significant transition rate × task interaction.

This suggests a possible sex-specific divergence in generalizability of gaze measures of spatial cognition. Together, these patterns raise the possibility that domain-general gaze-performance relationships may be more strongly expressed in males, whereas female gaze-performance relationships may be more sensitive to the specific demands of the task. However, larger sample sizes and direct tests of sex moderation are required to statistically validate these observations as evidence of sex-dependent gaze behaviour differences.

### 4.3 Within-task sex differences

#### 4.3.1 Sex differences in task performance

Exploratory within-task analyses also revealed several patterns of sex divergence in spatial processing. The Q-bitz *totalcorrect* difficulty × sex interaction showed a pattern of greater female decline in performance from P to R that did not survive correction, suggesting a greater rotation cost. Q-bitz RnP regression analyses showed a greater number of significant gaze predictors of performance in males (four) than females (one); while this could be due to overfitting of the exploratory analyses, it may also align with the more consistent, task-general gaze behaviour observed in males. While no significant sex differences were found in BBT or MRT raw score (*MRTscore*), a significant sex difference was found in MRT accuracy (*MRT%score*) with males scoring more accurately than females. Inspection of individual performance suggested that this difference was influenced by several female participants who struggled with the MRT and completed the task quickly and with low accuracy.

#### 4.3.2 Sex differences in Q-bitz gaze behaviour

The Q-bitz *fixationdurationcubes* ANOVA showed an overall greater fixation duration to cubes in males compared to females, and the interaction showed a decrease in females from P to R but no decrease in males. While neither effect survived multiple-comparison correction, this may reflect a greater gaze reallocation to the source in females, aligning with the well-documented male advantage in mental rotation (Voyer et al., 1995; Peters et al., 1995). This may also suggest that the difficulty-modulated changes in resource inspection found in the present study were more strongly expressed in females. A main effect of sex which survived correction was seen in Q-bitz *fixationdurationreplication*, in which males fixated longer to the replication. Like resource fixation, this may reflect differences in proportional gaze allocation to the source as task demands increase. *AOItransitionscard* was also higher in females than males. Though this did not survive correction, it may similarly reflect greater sampling of source information in females.

#### 4.3.3 Sex differences in BBT gaze behaviour

For the BBT, main effects of sex were seen in *pupildiameter* and *fixationdurationallpieces*. Females had greater pupil diameters than males, suggesting greater cognitive effort or load in female participants. In *fixationdurationallpieces*, longer fixation durations to the building supply were observed in males but did not survive correction. This pattern may reflect the established female advantage in perceptual speed (Majeres, 1977; Roivainen, 2011), which could allow for faster identification of the correct brick. Analyses revealed an interaction between difficulty × sex due to a significant increase in *fixationdurationreplication* from LMR to HMR which was present in females but not in males. Female participants fixated longer on their replication during the HMR condition when compared to the LMR, which may reflect greater inspection demands when navigating the 3-dimensional complexity of the HMR structure. A difficulty × sex interaction was also observed in *AOItransitionstotal*, where female transition rates were higher in LMR, but both sexes were similar in HMR. While it did not survive correction, it aligns with our findings on transition rate across tasks; the greater female decrease in transition rate from LMR to HMR may indicate greater relative increase in task demands for females. This interpretation is consistent with the greater sensitivity of female gaze behaviour to task demands observed across analyses.

#### 4.3.4 Sex differences in MRT gaze behaviour

A trial outcome × sex interaction was seen in MRT *pupildiameter* which did not survive correction, where female pupil diameter was significantly larger in incorrect trials than correct trials but male pupil diameter had no significant difference. Like in the BBT *pupildiameter* main effect of sex, this may suggest a higher relative cognitive load in female participants on trials resulting in incorrect responses. An interaction was also observed in MRT *fixationdurationtruecorrect* which did not survive correction, in which females fixated significantly longer to true correct answers in incorrect trials than correct trials, and males did not. While this does not align with any task-general gaze findings, it may be related to the higher degree of MRT accuracy in males. One possibility is that longer inspection of the true correct answers in females reflects greater uncertainty or difficulty resolving the correct response on trials that ultimately resulted in an error.

### 4.4 Limitations

While the present study has reinforced the validity of eye tracking in naturalistic tasks and the presence of domain-general gaze behaviours in spatial cognition, important limitations were encountered. Most importantly, future studies should include more consistent measures of both performance and gaze behaviour for more robust cross-task comparison.

Because BBT performance was measured by time, circularity was introduced in predictive analyses with time-dependent variables such as *AOItransitions*. As a result, the BBT was partly or completely excluded from many analyses such as within-task regressions and the transition rate mixed effects model. Future studies should implement time restrictions and use the number of bricks placed to assess performance, mirroring the present study’s Q-bitz task.

The MRT could not be directly compared in difficulty analyses due to an absence of difficulty conditions. Correct and incorrect trials were instead compared as a function of trial outcome. Although incorrect trials may reflect instances in which task demands exceeded successful spatial processing, they cannot be considered equivalent to experimentally manipulated levels of difficulty. In the future, MRT trials should be categorized by degree of rotation, which corresponds with trial difficulty (Shepard & Metzler, 1971). To account for participants completing a different number of trials within the three-minute time limit, trials should be administered in alternating order of easy/hard, so each participant completes the same proportion of both difficulty conditions (+/− one trial for odd-numbers of completion).

Source inspection and transition rate could be directly compared across all tasks, but other gaze behaviours could not be due to differing areas of interest. The MRT did not have an equivalent area for resource inspection, while Q-bitz and BBT did not have equivalent areas for MRT *truecorrect* and *trueincorrect*. Additionally, gaze behaviours involving the replication space (and equivalently, MRT answers) were not consistent due to differing demands of each task. For example, the replication demanded greater motor coordination to complete in BBT than Q-bitz.

Not all eye tracking measures could be included in regressions due to overlap leading to multicollinearity. For example, only one of the fixation proportion measures was included for each task. Excluded measures may nonetheless be related to performance and therefore may represent significant predictors which were not detected.

Pupillometry was framed exploratorily in the present study due to difficult isolation of cognitive load mediated pupil dilation in a naturalistic setting. While a significant effect of *pupildiameter* was observed in BBT, the MRT effect did not survive correction, and none were found in Q-bitz; inconsistent with the presence of other measures indicative of cognitive load differences. Specific confounding factors observed included anticipatory pupil dilation in the normalization period and fatigue-mediated pupil constriction across longer tasks like the BBT and MRT.

Saccade amplitude significance was only observed in the BBT; as a main effect of difficulty (greater in LMR) and as a negative predictor of performance in the LMR condition (smaller saccade amplitudes predicted higher time). Although this is consistent with established association of smaller saccade amplitudes with increased difficulty (May et al., 1990), it did not arise in the other two tasks. This is likely due to the fixed nature of the Q-bitz and MRT, where gaze is confined to several specific areas, thus decreasing variation in saccade amplitude. While saccade amplitude was not an effective measure for the present study, the BBT results suggest that it may be more sensitive to the greater visual search, spatial extent, or visuomotor demands of this task, although the present design cannot determine which task characteristic produced this effect.

Because the present study was unable to fully utilize pupillometry, cognitive load could only be inferred (rather than directly measured) through other eye tracking measures. Although cognitive load has been linked to difficulty (Paas & Van Merriënboer, 1993) and various eye tracking measures, these relationships are not causative and may be confounded by other factors beyond cognitive effort (Paas & Van Merriënboer, 1993), such as engagement and motor demands. However, each significant eye tracking measure in the present study independently varies in the direction that supports its link to cognitive load. Specifically, where significant, pupil diameter increases with load (Bauer et al., 2022), saccade amplitude decreases with load (May et al., 1990), fixation duration increases with load (Just & Carpenter, 1980), and transition rate decreases with load (Recarte & Nunes, 2003). The convergence of these individual measures strengthens the consistency of non-pupillometry measures with cognitive load. Future studies could combine eye tracking with an independent measure of cognitive load to directly test whether the observed gaze changes correspond to changes in cognitive effort.

Despite gaze behaviours suggesting increased cognitive load in females across all tasks, many of the sex differences in gaze behaviour did not survive multiple-comparison correction. Additionally, both BBT *time* and MRT raw score (*MRTscore*) failed to replicate established findings of significant male performance advantages (Aguilar Ramirez et al., 2020; 2021; 2022). Although *MRTscore* did not produce sex differences, the increased female difficulty was significantly reflected in *MRT%score*. This was likely due to the limited size of the sample (*N* = 30) as well as the decreased scope of tasks compared to previous studies. While the current study used one set of 12 trials for the MRT and one model from each BBT condition, previous studies have used two sets of MRT trials (for a total of 24 questions in six minutes) and up to three models from each BBT condition (Aguilar Ramirez et al., 2020; 2021; 2022). Additionally, the sex-separated cross-task analyses were exploratory and did not directly test whether gaze-performance relationships differed significantly between males and females. Future work with larger samples and direct tests of sex moderation will be needed to determine whether the sex-specific patterns observed in the present study reflect reliable differences in gaze behaviour.

### 4.5 Conclusion

In conclusion, the present study assessed the preservation and generalizability of gaze behaviours across naturalistic spatial tasks with different perceptual, cognitive and visuomotor demands. An increase in source inspection and decrease in transition rates was observed with increased task demands and poorer task performance, consistent with decreased encoding efficiency and increased cognitive load. Source fixation duration was shown to be a task-invariant predictor of performance. In other words, the relationship between source fixation duration and task performance did not significantly differ across tasks, identifying source fixation duration as a domain-general predictor of spatial performance. This suggests that efficient encoding of source information may represent a common cognitive process underlying performance across different spatial tasks. While transition rate also predicted performance across tasks, the strength of the relationship was not consistent. This suggests that although transition rate may represent a stable gaze tendency across tasks, its relationship with performance is more strongly modulated by task-specific demands.

Within each task, a variety of eye tracking metrics were revealed to predict performance and differ between difficulty conditions. While the directionality of all associations between gaze, performance, and difficulty was consistent with previous work, different significant metrics arose in each task. Notably, significant differences in saccade amplitude were consistently seen in the BBT but no other task; while significant differences in pupil diameter mostly arose in the MRT. This suggests that although gaze behaviour can reflect cognitive processes across naturalistic spatial tasks, the sensitivity of individual gaze metrics may depend on the specific perceptual, cognitive, and visuomotor demands of the task.

Sex-specific exploratory analyses also revealed patterns suggesting potential sex differences in the generalizability of gaze behaviours during spatial solving. While this aligns with established sex differences in spatial ability, larger samples and direct tests of sex moderation are needed to determine whether these patterns reflect reliable sex differences in gaze-performance relationships.

Overall, this study demonstrated the utility of eye tracking for assessing spatial cognition across naturalistic tasks. It highlighted the differing strengths and limitations of specific eye tracking measures across task contexts and established the need for further assessment of sex differences.

## Appendix A

### Q-bitz Correlations

**Figure A1:**
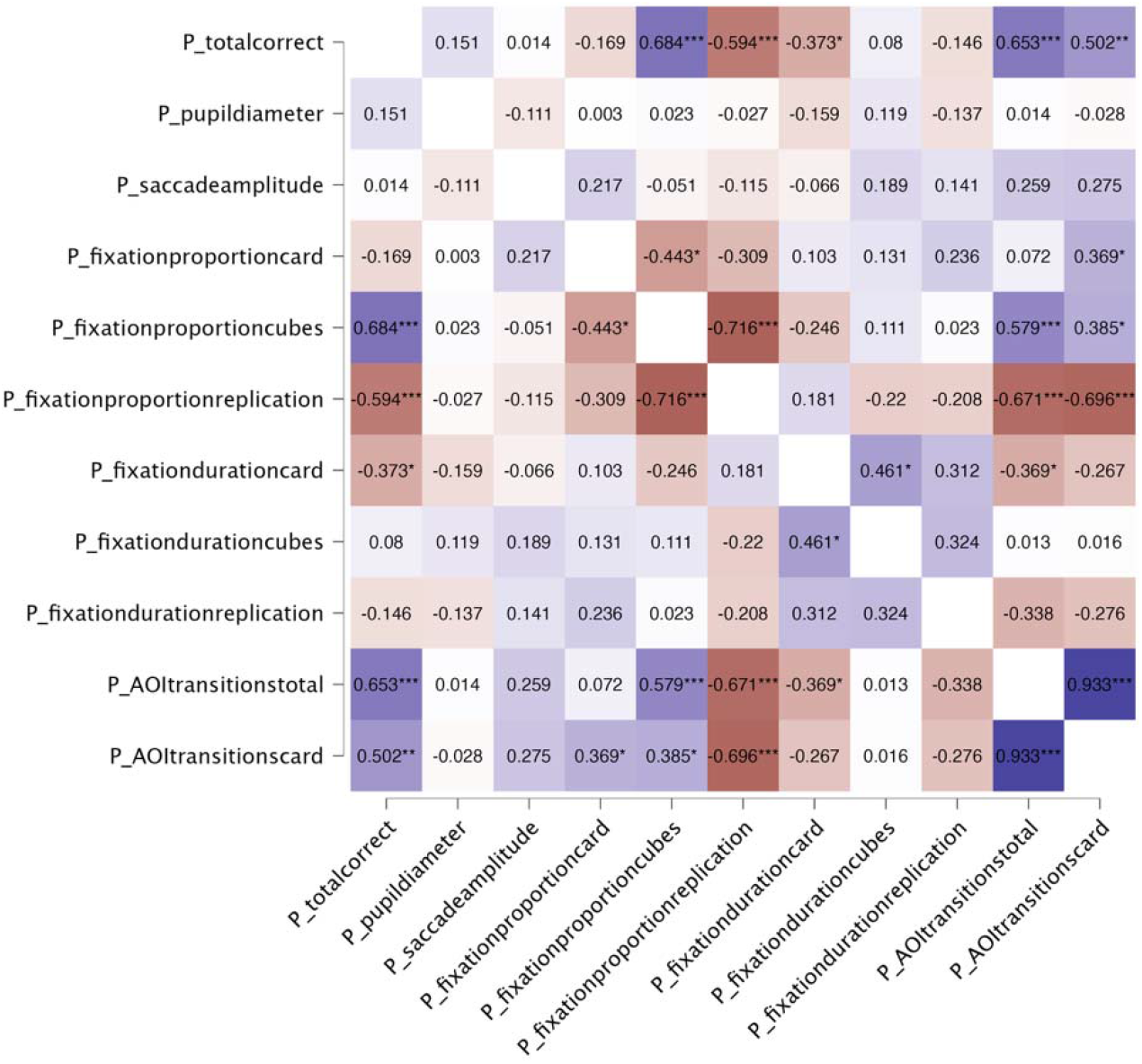
correlations of all measures in Q-bitz Pattern (P) condition for all participants. Significant correlations are marked with asterisks (* p < .05, ** p < .01, *** p < .001). Positive correlations are shown in blue, negative correlations in red, with colour intensity proportional to magnitude of the coefficient.

In the P condition with all participants, *totalcorrect* is positively correlated with *fixationproportioncubes* (*r* (28) = .68, *p* < .001), *AOItransitionstotal* (*r* (28) = .65, *p* < .001), and *AOItransitionscard* (*r* (28) = .50, *p* = .005), and negatively correlated with *fixationproportionreplication* (*r* (28) = –.59, *p* < .001) and *fixationdurationcard* (*r* (28) = – .37, *p* = .042).

**Figure A2:**
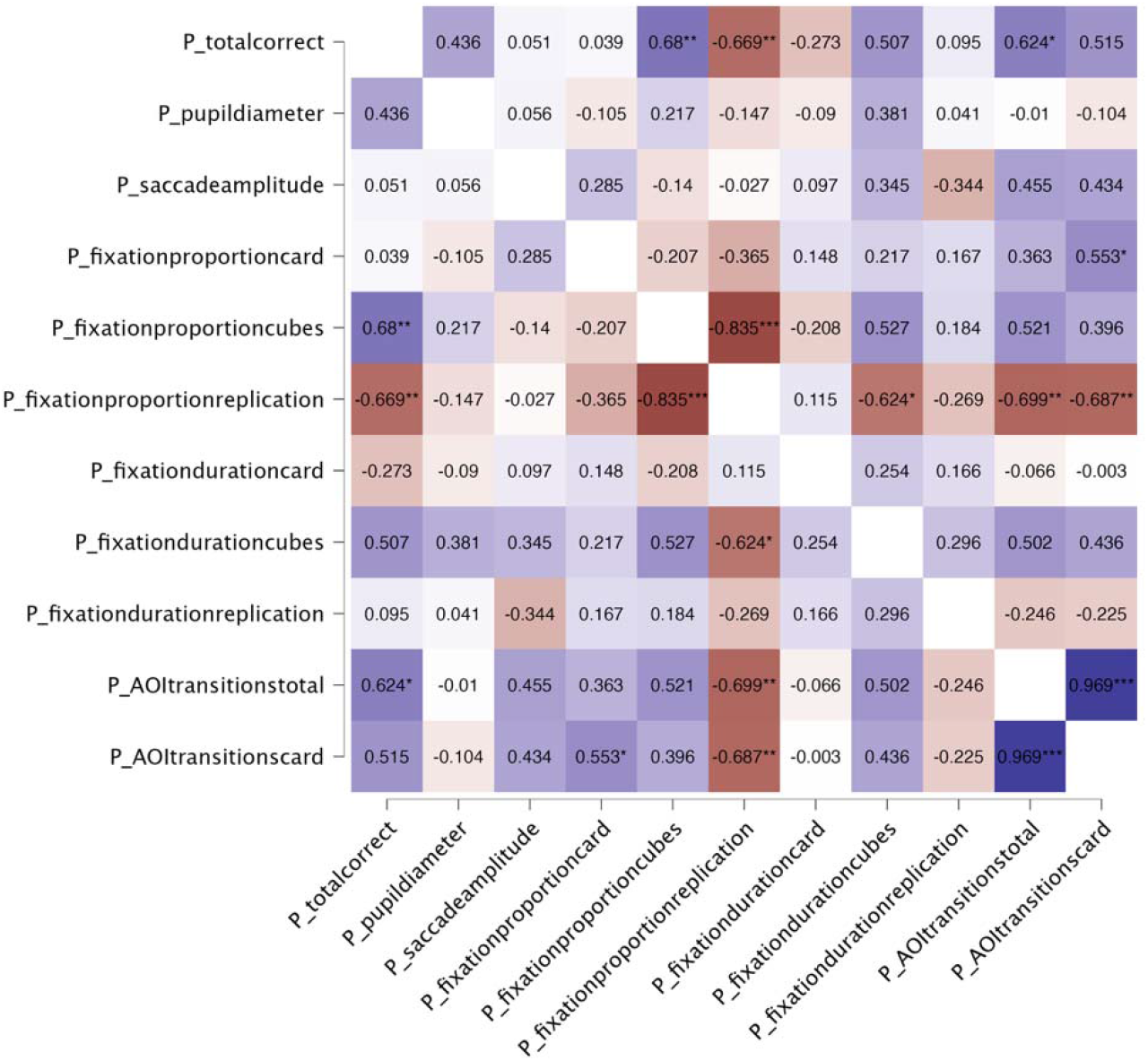
correlations of all measures in Q-bitz Pattern (P) condition for female participants. Significant correlations are marked with asterisks (* p < .05, ** p < .01, *** p < .001). Positive correlations are shown in blue, negative correlations in red, with colour intensity proportional to magnitude of the coefficient.

In the P condition with female participants only, *totalcorrect* is positively correlated with *fixationproportioncubes* (*r* (12) = .68, *p* = .007) and *AOItransitionstotal* (*r* (12) = .62, *p* = .017) and approaches positive correlation with *fixationdurationcubes* (*r* (12) = .51, *p* = .064) and *AOItransitionscard* (*r* (12) = .515, *p* = .059). It is also negatively correlated with *fixationproportionreplication* (*r* (12) = –.67, *p* = .009).

**Figure A3:**
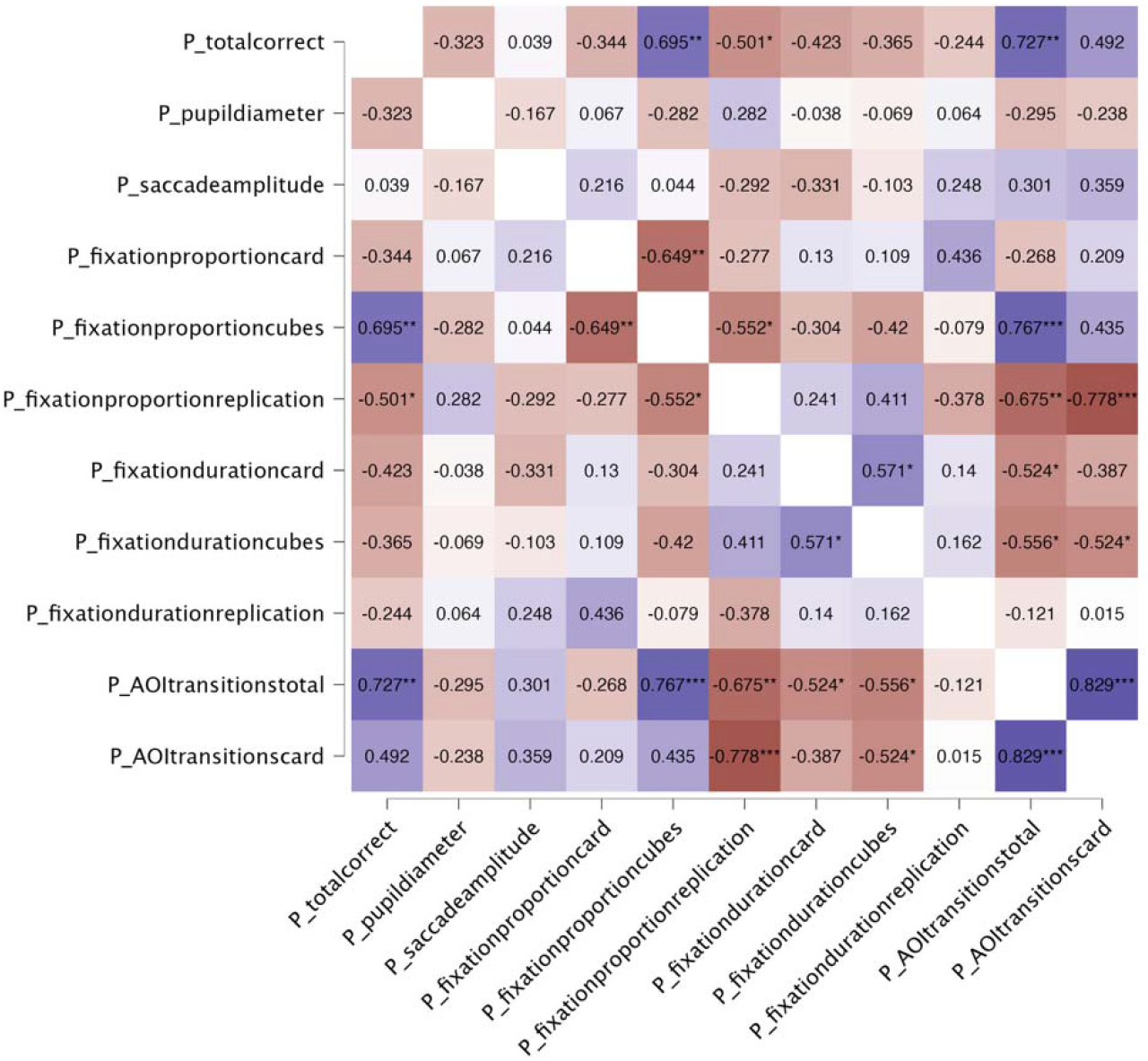
correlations of all measures in Q-bitz Pattern (P) condition for male participants. Significant correlations are marked with asterisks (* p < .05, ** p < .01, *** p < .001). Positive correlations are shown in blue, negative correlations in red, with colour intensity proportional to magnitude of the coefficient.

In the P condition with male participants only, *totalcorrect* is positively correlated with *fixationproportioncubes* (*r* (14) = .70, *p* = .003) and *AOItransitionstotal* (*r* (14) = .73, *p* = .001) and approaches positive correlation with *AOItransitionscard* (*r* (14) = .49, *p* = .053). It is also negatively correlated with *fixationproportionreplication* (*r* (14) = –.50, *p* = .048).

**Figure A4:**
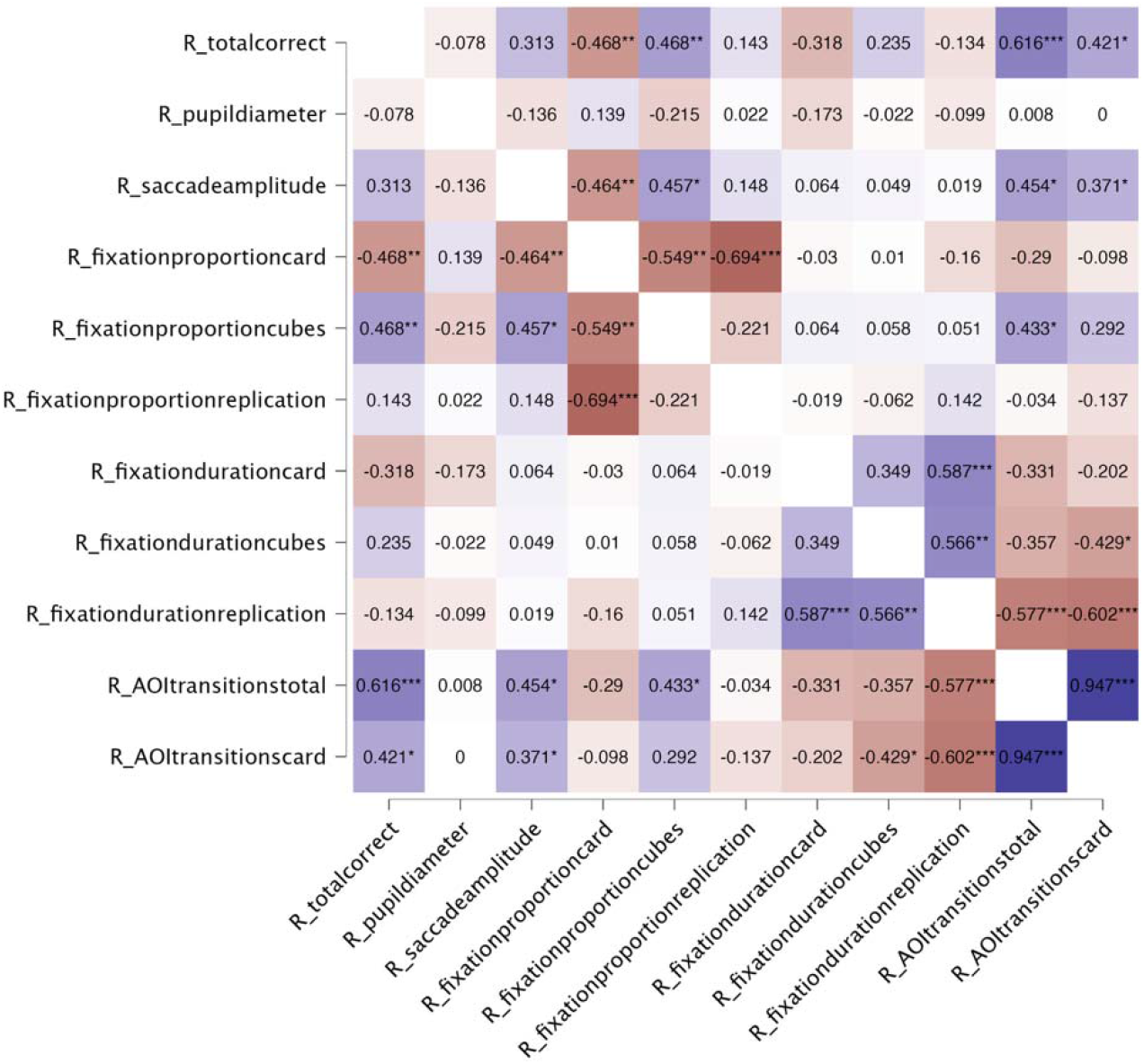
correlations of all measures in Q-bitz Rotated pattern (R) condition for all participants. Significant correlations are marked with asterisks (* p < .05, ** p < .01, *** p < .001). Positive correlations are shown in blue, negative correlations in red, with colour intensity proportional to magnitude of the coefficient.

In the R condition with all participants, *totalcorrect* is positively correlated with *fixationproportioncubes* (*r* (28) = .47, *p* = .009), *AOItransitionstotal* (*r* (28) = .62, *p* < .001), and *AOItransitionscard* (*r* (28) = .42, *p* = .021), and approaches positive correlation with *saccadeamplitude* (*r* (28) = .31, *p* = .092). It is also negatively correlated with *fixationproportioncard* (*r* (28) = –.47, *p* = .009) and approaches negative correlation with *fixationdurationcard* (*r* (28) = –.32, *p* = .087).

**Figure A5:**
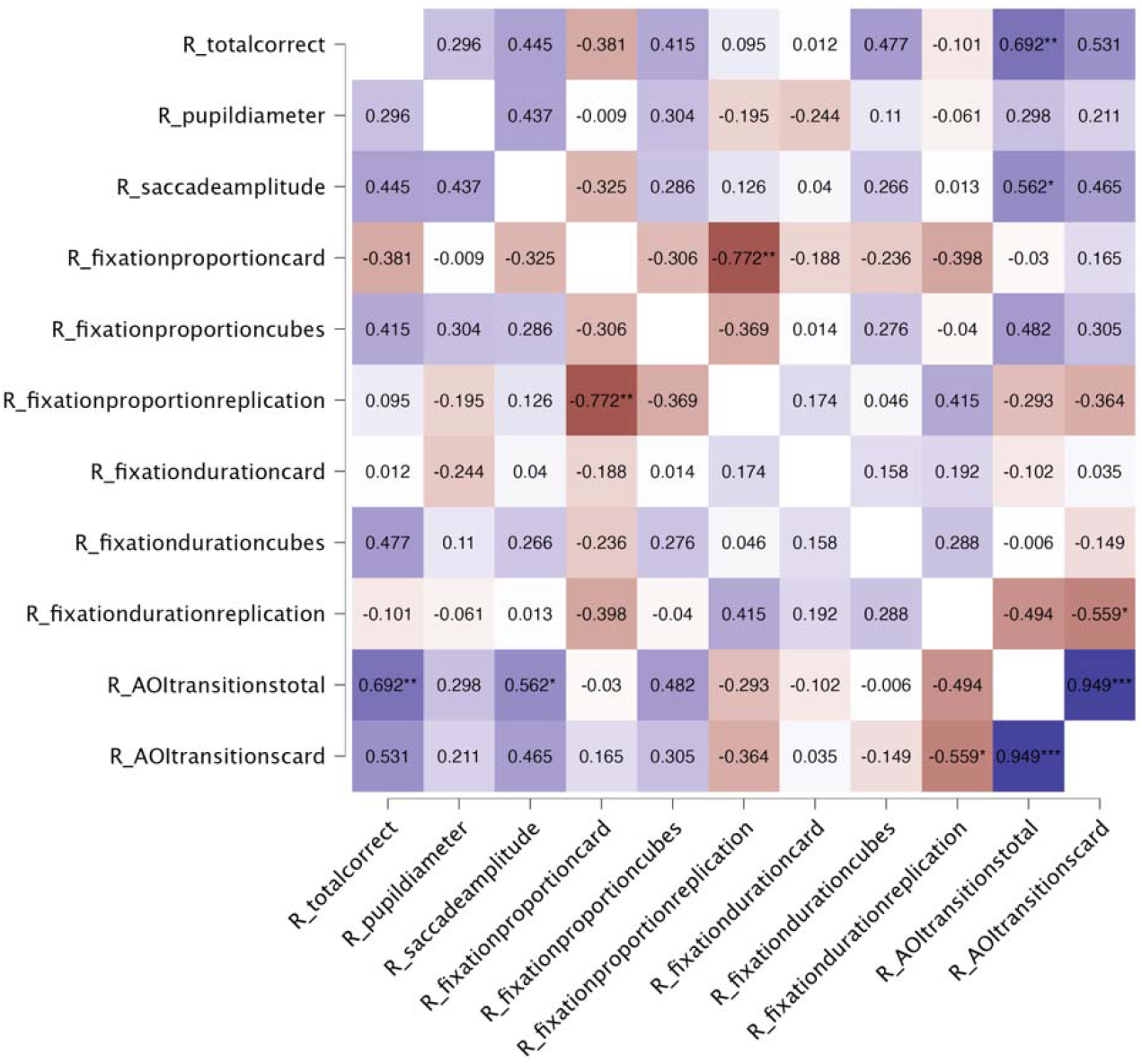
correlations of all measures in Q-bitz Rotated pattern (R) condition for female participants. Significant correlations are marked with asterisks (* p < .05, ** p < .01, *** p < .001). Positive correlations are shown in blue, negative correlations in red, with colour intensity proportional to magnitude of the coefficient.

In the R condition with female participants only, *totalcorrect* is positively correlated with *AOItransitionstotal* (*r* (12) = .69, *p* = .006) and approaches positive correlation with *fixationdurationcubes* (*r* (12) = .48, *p* = .085) and *AOItransitionscard* (*r* (12) = .53, *p* = .051). No negative correlations were present.

**Figure A6:**
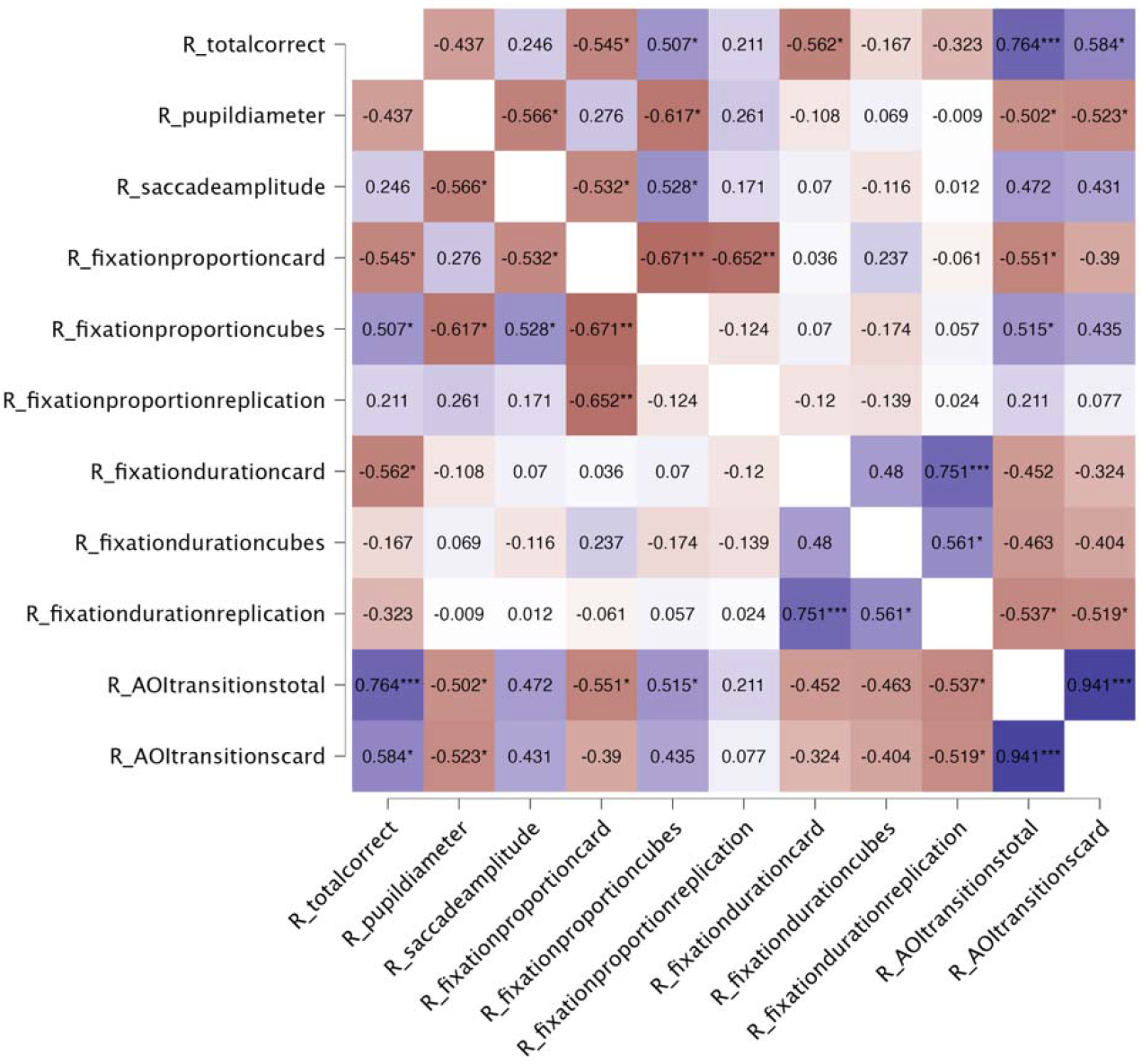
correlations of all measures in Q-bitz Rotated pattern (R) condition for male participants. Significant correlations are marked with asterisks (* p < .05, ** p < .01, *** p < .001). Positive correlations are shown in blue, negative correlations in red, with colour intensity proportional to magnitude of the coefficient.

In the R condition with male participants only, totalcorrect is positively correlated with fixationproportioncubes (*r* (14) = .51, *p* = .045), AOItransitionstotal (*r* (14) = .76, *p* < .001), and AOItransitionscard (*r* (14) = .58, *p* = .018). It is negatively correlated with fixationproportioncard (*r* (14) = –.55, *p* = .029) and fixationdurationcard (*r* (14) = –.56, *p* = .023) and approaches negative correlation with pupildiameter (*r* (14) = –.44, *p* = .091).

**Figure A7:**
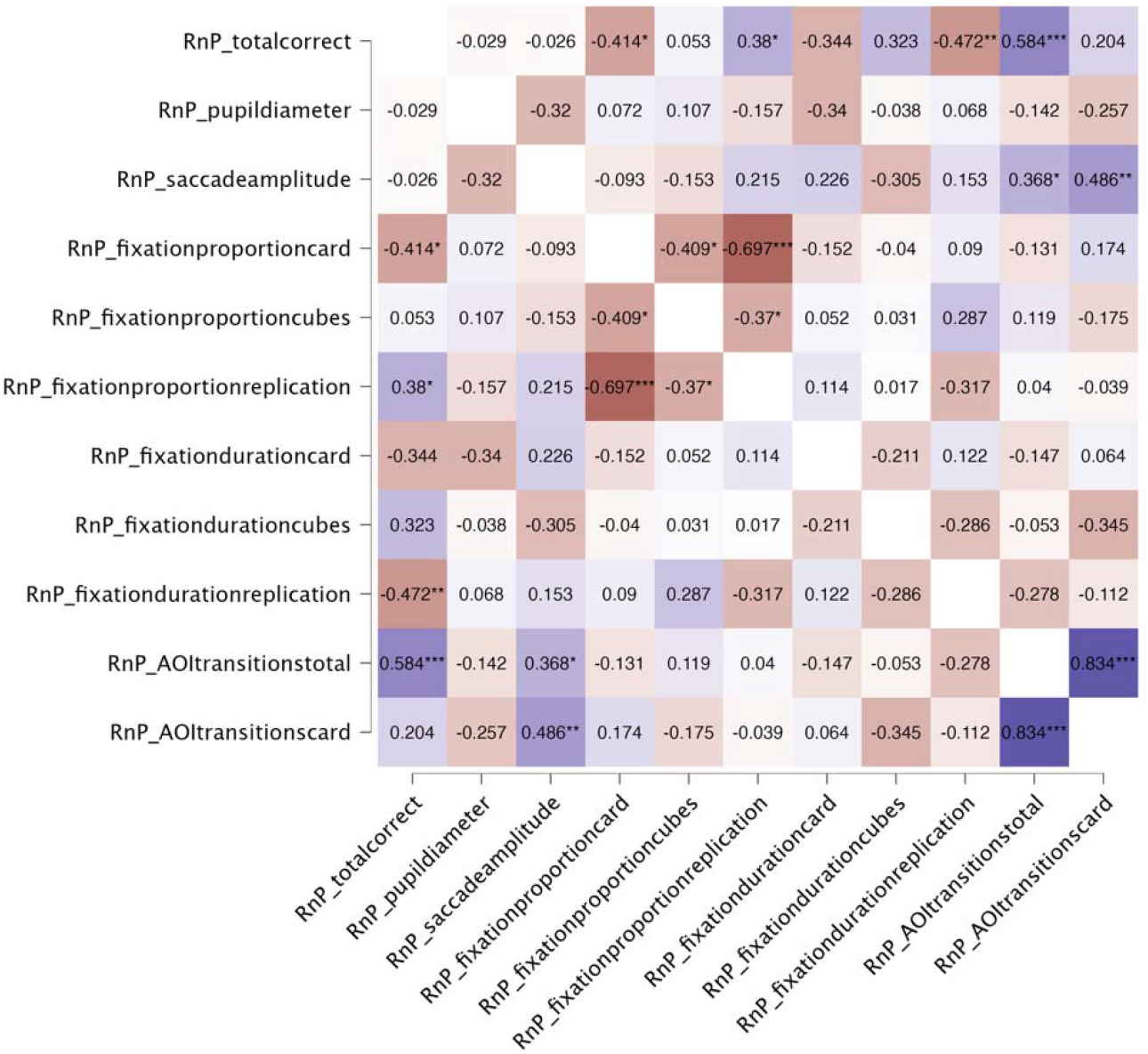
correlations of all measures in Q-bitz R normalized to P condition for all participants. Significant correlations are marked with asterisks (* p < .05, ** p < .01, *** p < .001). Positive correlations are shown in blue, negative correlations in red, with colour intensity proportional to magnitude of the coefficient.

With all participants included, totalcorrect is positively correlated with *fixationproportionreplication* (*r* (28) = .38, *p* = .038) and *AOItransitionstotal* (*r* (28) = .58, *p* < .001) and approaches positive correlation with *fixationdurationcubes* (*r* (28) = .32, *p* = .081). It is also negatively correlated with *fixationproportioncard* (*r* (28) = –.41, *p* = .023) and *fixationdurationreplication* (*r* (28) = –.47, *p* = .008) and approaches negative correlation with *fixationdurationcard* (*r* (28) = –.34, *p* = .063).

**Figure A8:**
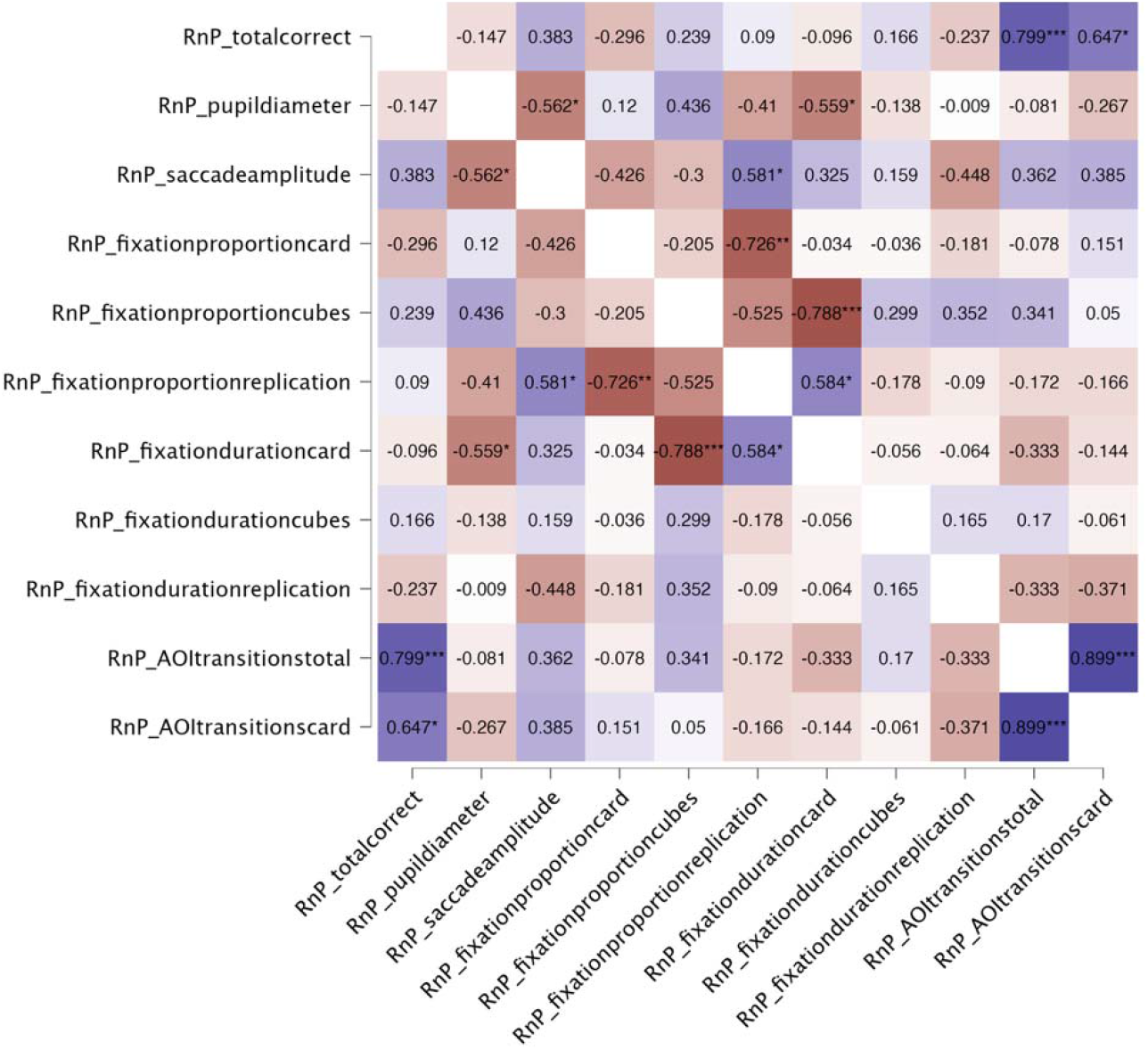
correlations of all measures in Q-bitz R normalized to P condition for female participants. Significant correlations are marked with asterisks (* p < .05, ** p < .01, *** p < .001). Positive correlations are shown in blue, negative correlations in red, with colour intensity proportional to magnitude of the coefficient.

With female participants only, *totalcorrect* is positively correlated with *AOItransitionstotal* (*r* (12) = .80, *p* < .001) and *AOItransitionscard* (*r* (12) = .65, *p* = .012). No negative correlations are present.

**Figure A9:**
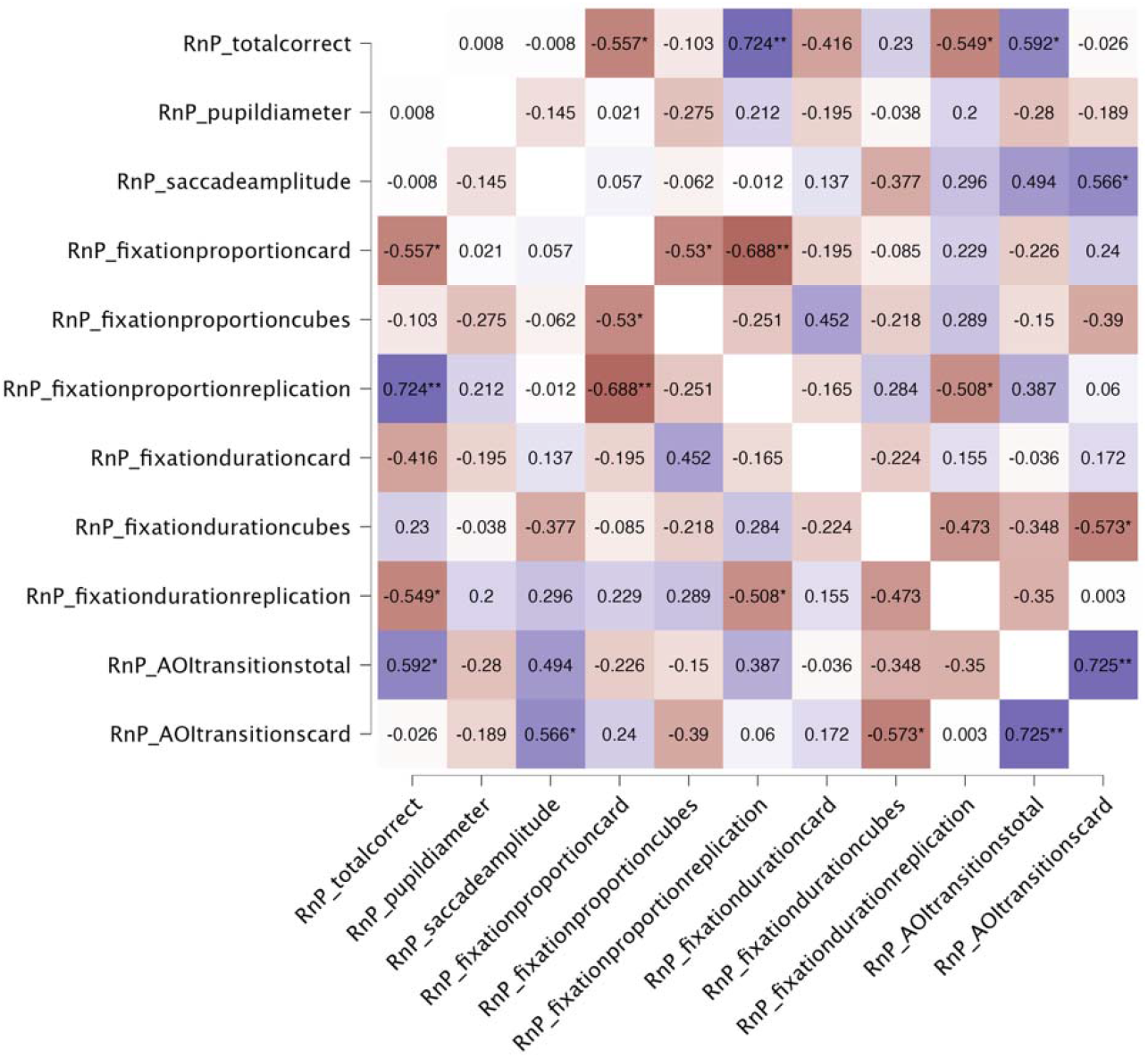
correlations of all measures in Q-bitz R normalized to P condition for male participants. Significant correlations are marked with asterisks (* p < .05, ** p < .01, *** p < .001). Positive correlations are shown in blue, negative correlations in red, with colour intensity proportional to magnitude of the coefficient.

With male participants only, *totalcorrect* is positively correlated with *fixationproportionreplication* (*r* (14) = .72, *p* = .002) and *AOItransitionstotal* (*r* (14) = .59, *p* = .016) and negatively correlated with *fixationproportioncard* (*r* (14) = –.56, *p* = .025) and *fixationdurationreplication* (*r* (14) = –.55, *p* = .028).

## Appendix B

### BBT Correlations

**Figure B1:**
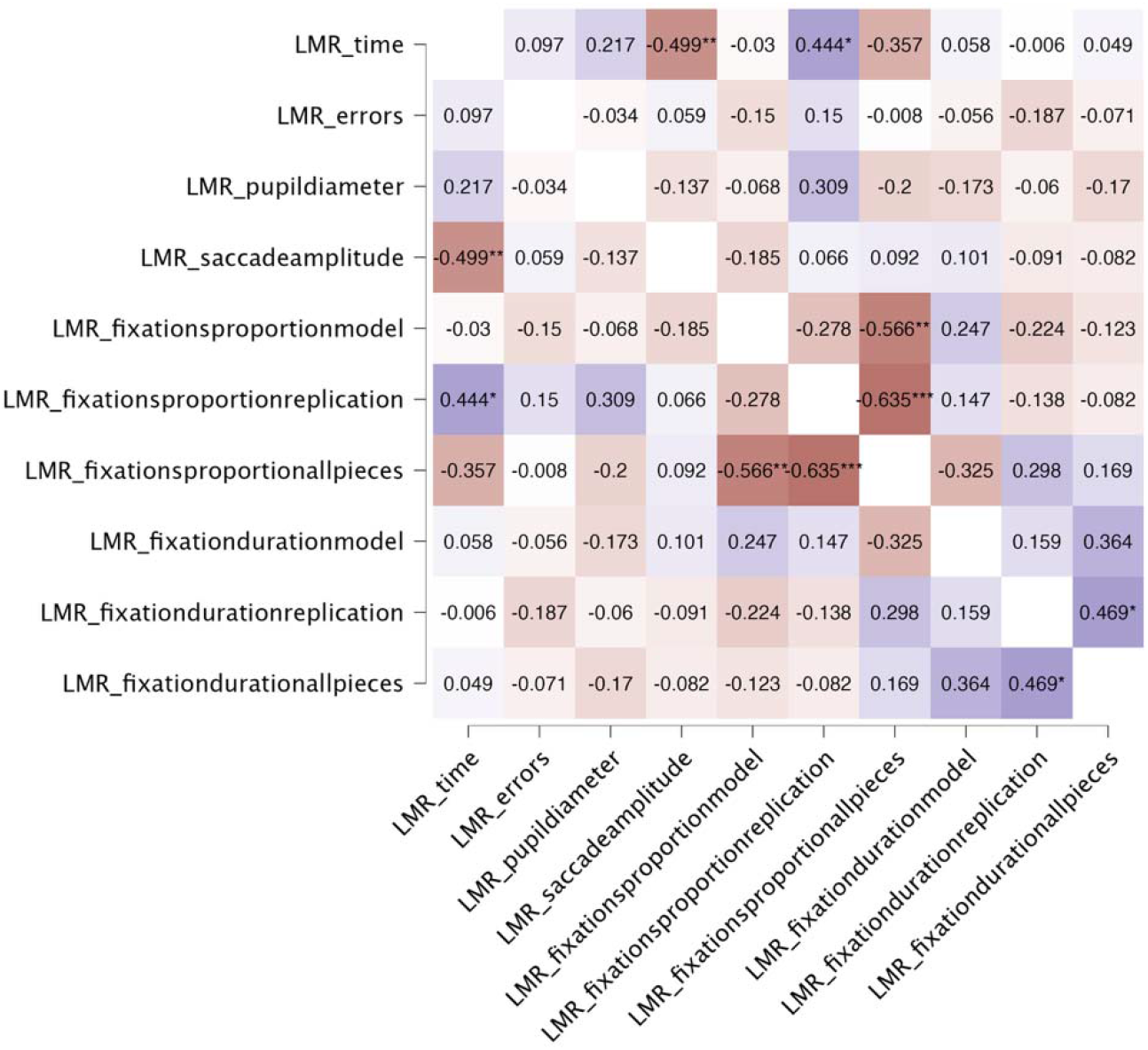
correlations of all measures in LMR condition for all participants. Significant correlations are marked with asterisks (* p < .05, ** p < .01, *** p < .001). Positive correlations are shown in blue, negative correlations in red, with colour intensity proportional to magnitude of the coefficient.

In the LMR condition with all participants, *time* is positively correlated with *fixationproportionreplication* (*r* (26) = .44, *p* = .018), negatively correlated with *saccadeamplitude* (*r* (28) = –.50, *p* = .005), and approaches negative correlation with *fixationproportionallpieces* (*r* (26) = –.36, *p* = .062).

**Figure B2:**
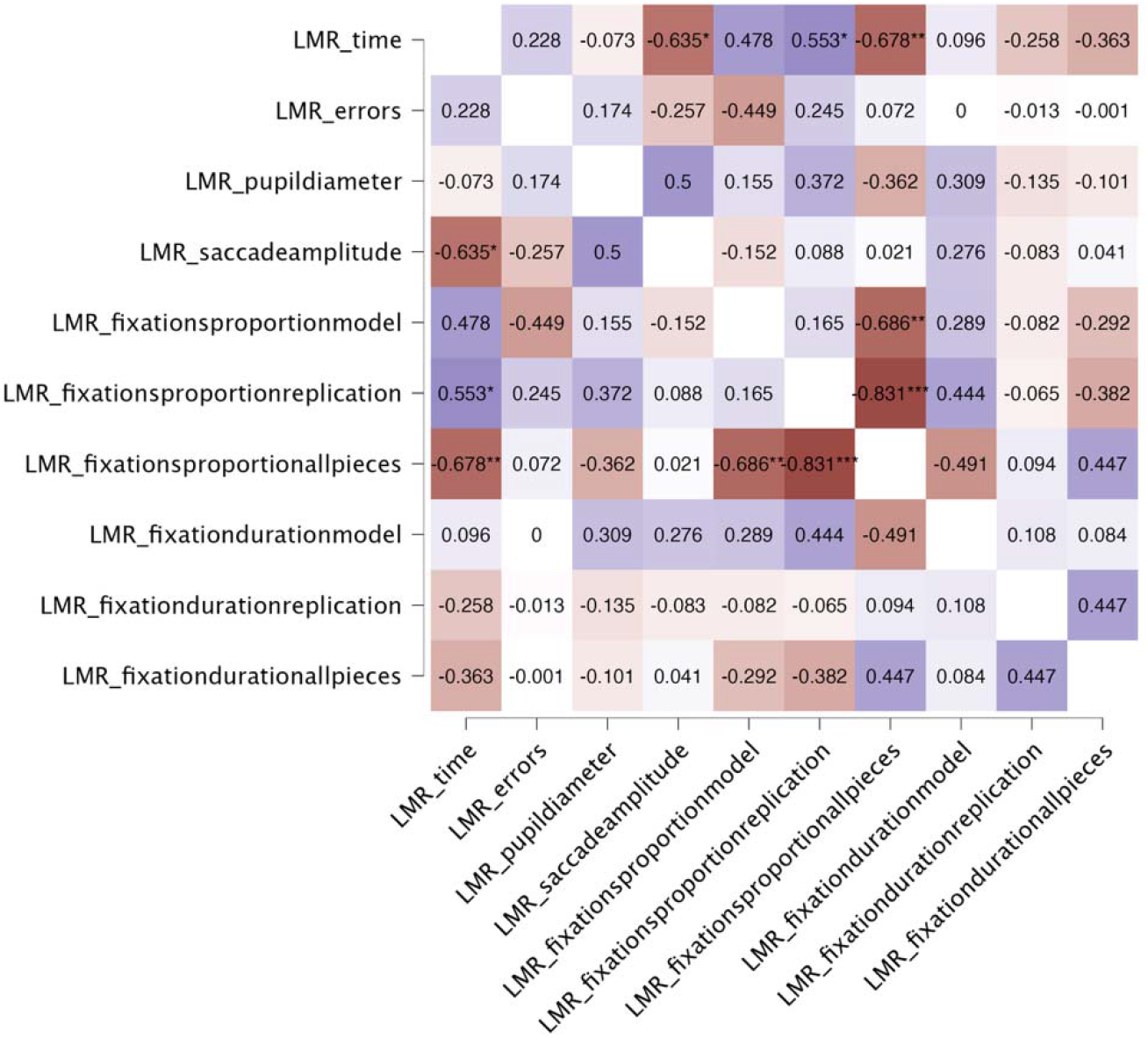
correlations of all measures in LMR condition for female participants. Significant correlations are marked with asterisks (* p < .05, ** p < .01, *** p < .001). Positive correlations are shown in blue, negative correlations in red, with colour intensity proportional to magnitude of the coefficient.

In the LMR condition with female participants only, *time* is positively correlated with *fixationproportionreplication* (*r* (12) = .55, *p* = .040) and approaches positive correlation with *fixationproportionmodel* (*r* (12) = .48, *p* = .084). It is also negatively correlated with *saccadeamplitude* (*r* (12) = –.64, *p* = .015) and *fixationproportionallpieces* (*r* (12) = –.68, *p* = .008).

**Figure B3:**
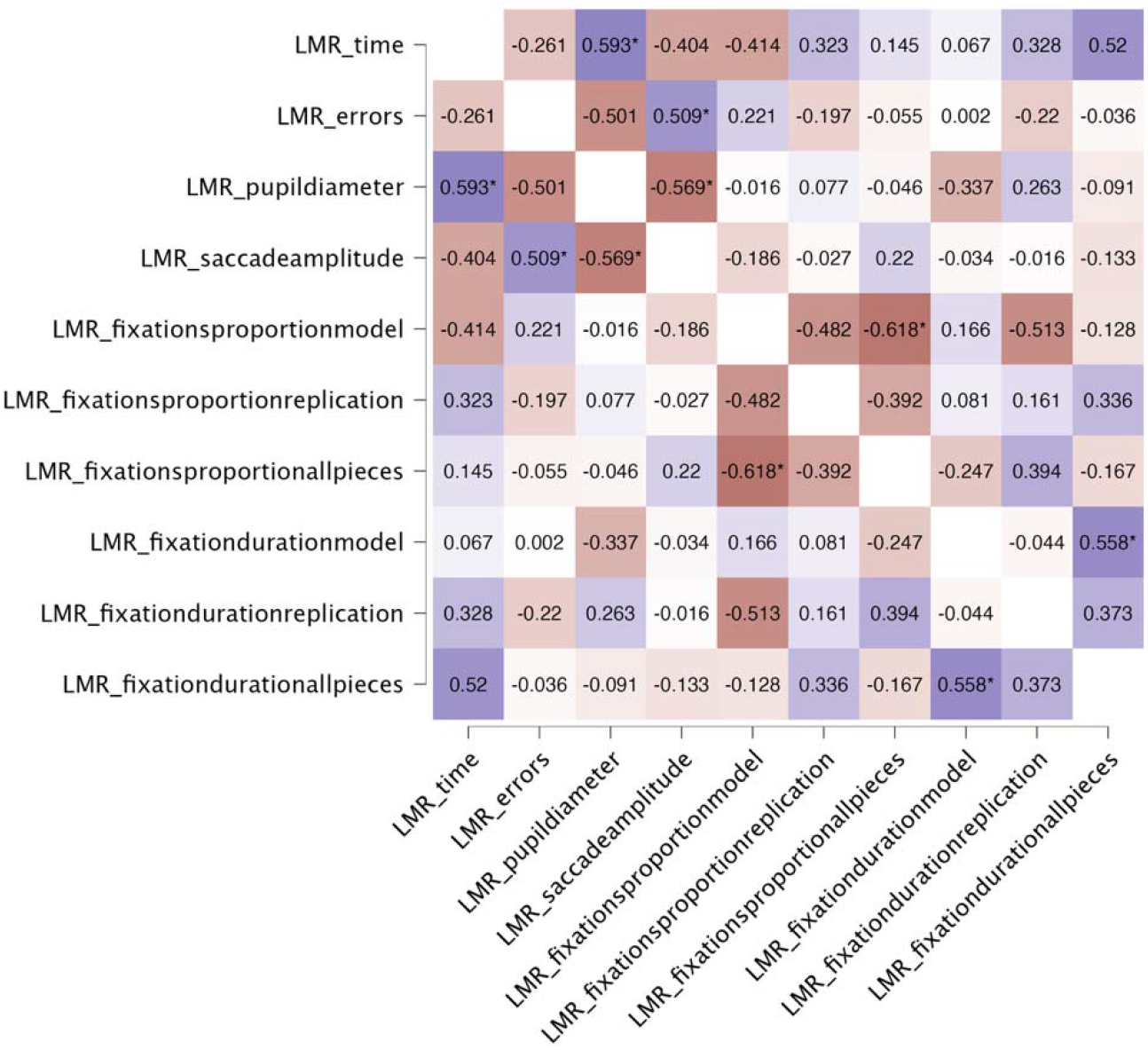
correlations of all measures in LMR condition for male participants. Significant correlations are marked with asterisks (* p < .05, ** p < .01, *** p < .001). Positive correlations are shown in blue, negative correlations in red, with colour intensity proportional to magnitude of the coefficient.

In the LMR condition with male participants only, *time* is positively correlated with *pupildiameter* (*r* (11) = .59, *p* = .033) and approaches positive correlation with *fixationdurationallpieces* (*r* (12) = .52, *p* = .057). No negative correlations are present.

**Figure B4:**
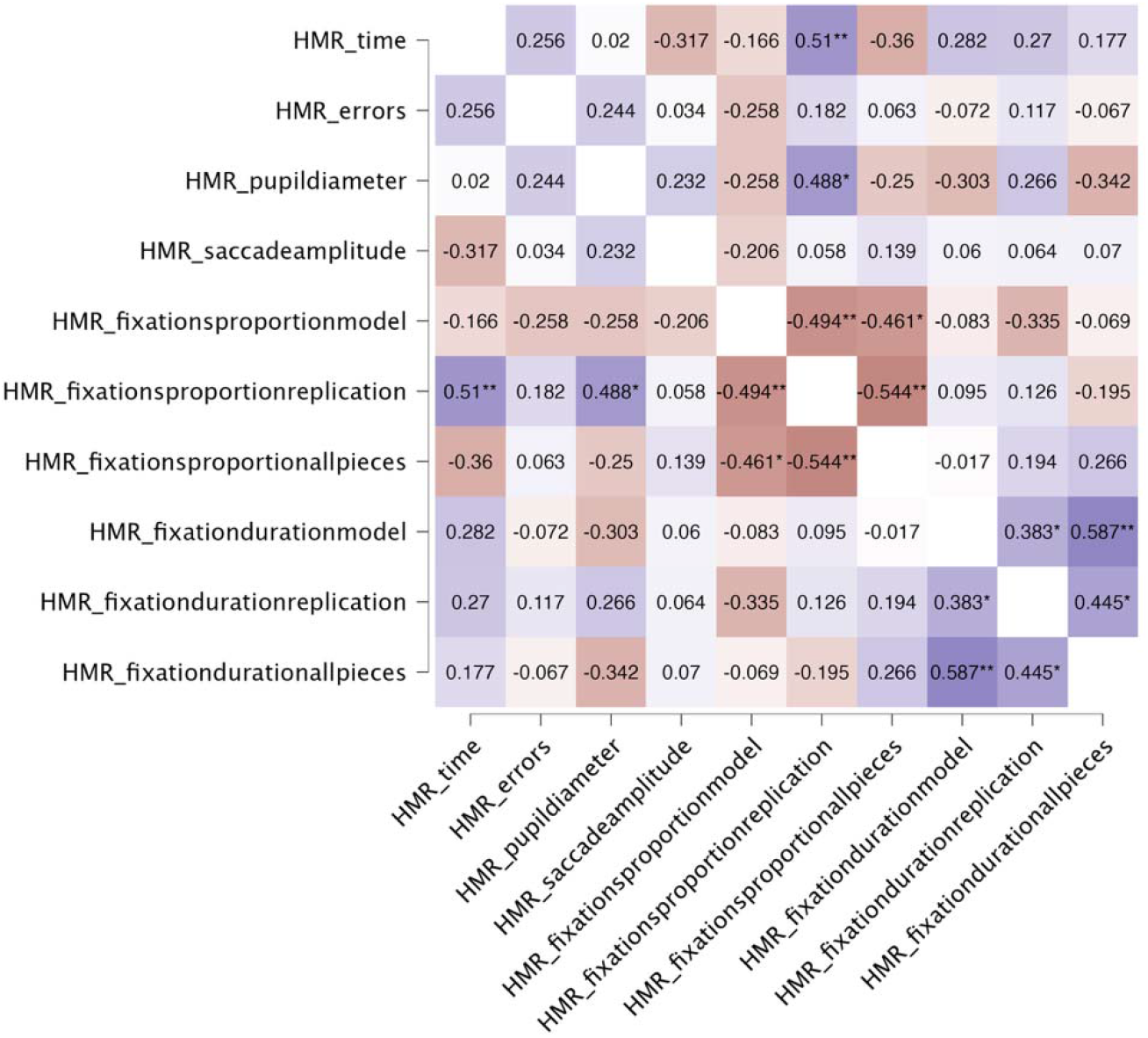
correlations of all measures in HMR condition for all participants. Significant correlations are marked with asterisks (* p < .05, ** p < .01, *** p < .001). Positive correlations are shown in blue, negative correlations in red, with colour intensity proportional to magnitude of the coefficient.

In the HMR condition with all participants, *time* is positively correlated with *fixationproportionreplication* (*r* (26) = .51, *p* = .006) and approaches negative correlation with *saccadeamplitude* (*r* (28) = –.32, *p* = .088) and *fixationproportionallpieces* (*r* (26) = – .36, *p* = .060).

**Figure B5:**
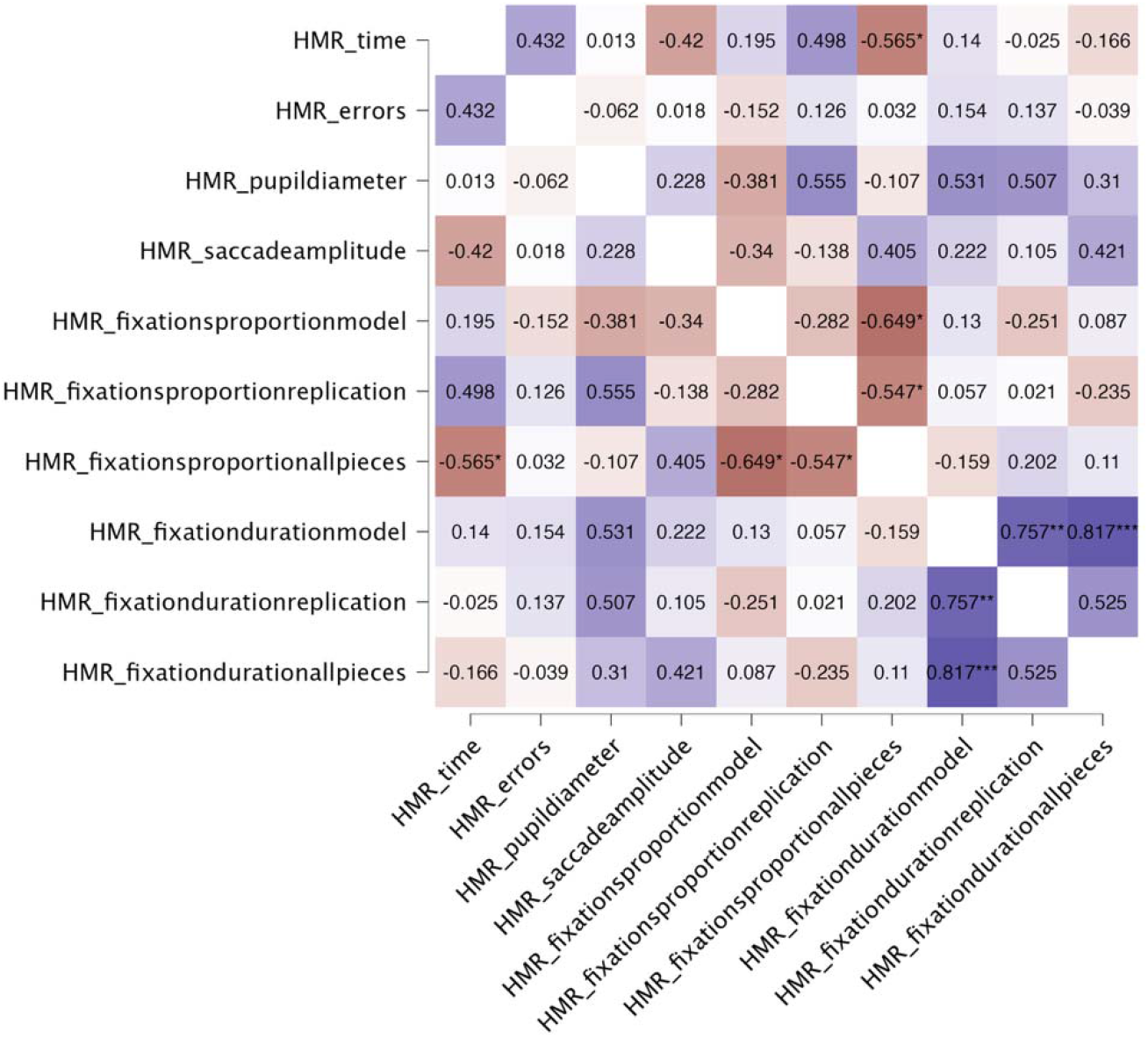
correlations of all measures in HMR condition for female participants. Significant correlations are marked with asterisks (* p < .05, ** p < .01, *** p < .001). Positive correlations are shown in blue, negative correlations in red, with colour intensity proportional to magnitude of the coefficient.

In the HMR condition with female participants only, *time* approaches positive correlation with *fixationproportionreplication* (*r* (12) = .50, *p* = .070) and is negatively correlated with *fixationproportionallpieces* (*r* (12) = –.57, *p* = .035).

**Figure B6:**
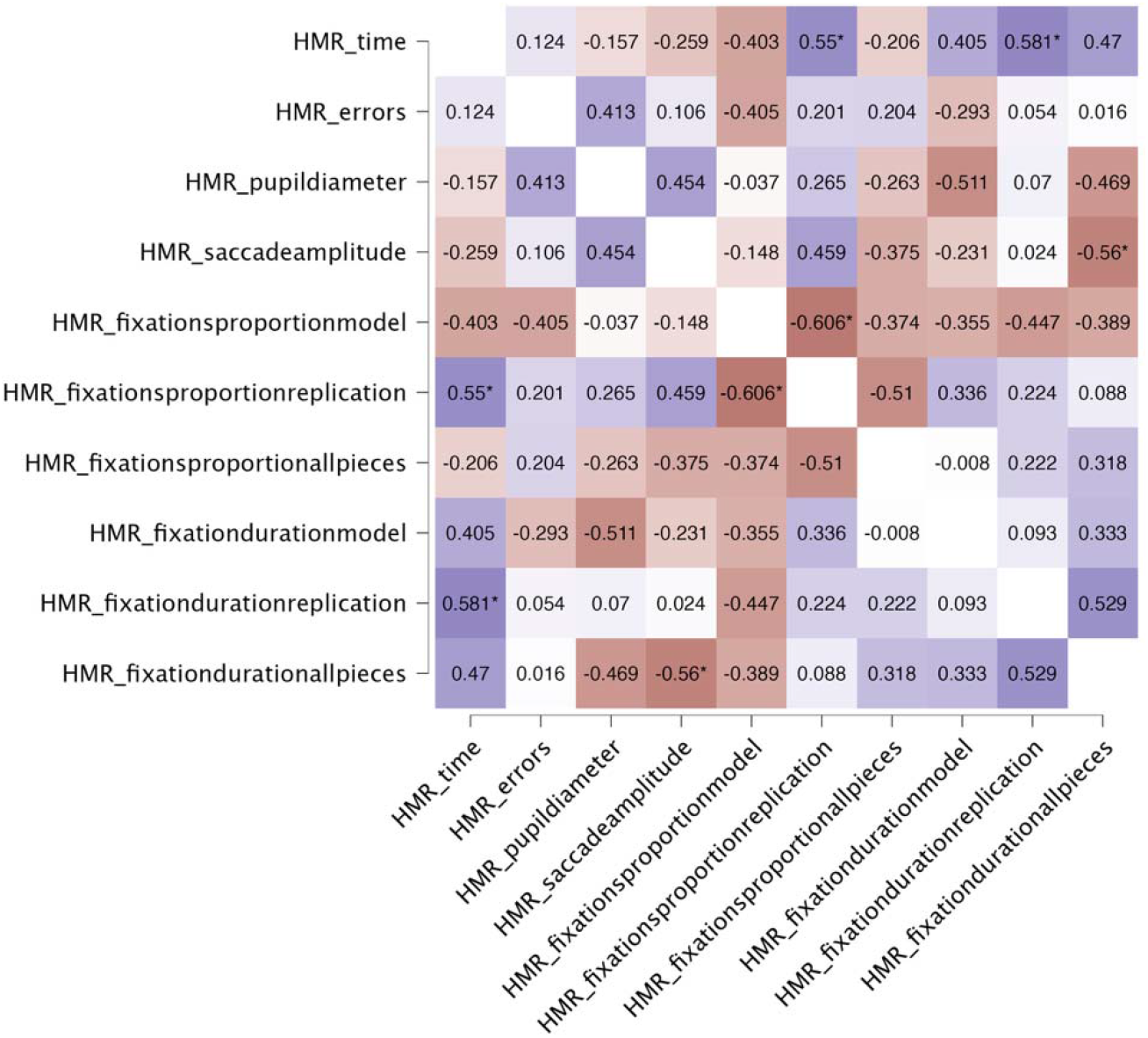
correlations of all measures in HMR condition for male participants. Significant correlations are marked with asterisks (* p < .05, ** p < .01, *** p < .001). Positive correlations are shown in blue, negative correlations in red, with colour intensity proportional to magnitude of the coefficient.

In the HMR condition with male participants only, *time* is positively correlated with *fixationproportionreplication* (*r* (12) = .55, *p* = .041) and *fixationdurationreplication* (*r* (12) = .58, *p* = .029) and approaches positive correlation with *fixationdurationallpieces* (*r* (12) = .47, *p* = .090). No negative correlations are present.

## Appendix C

### MRT Correlations

**Figure C1:**
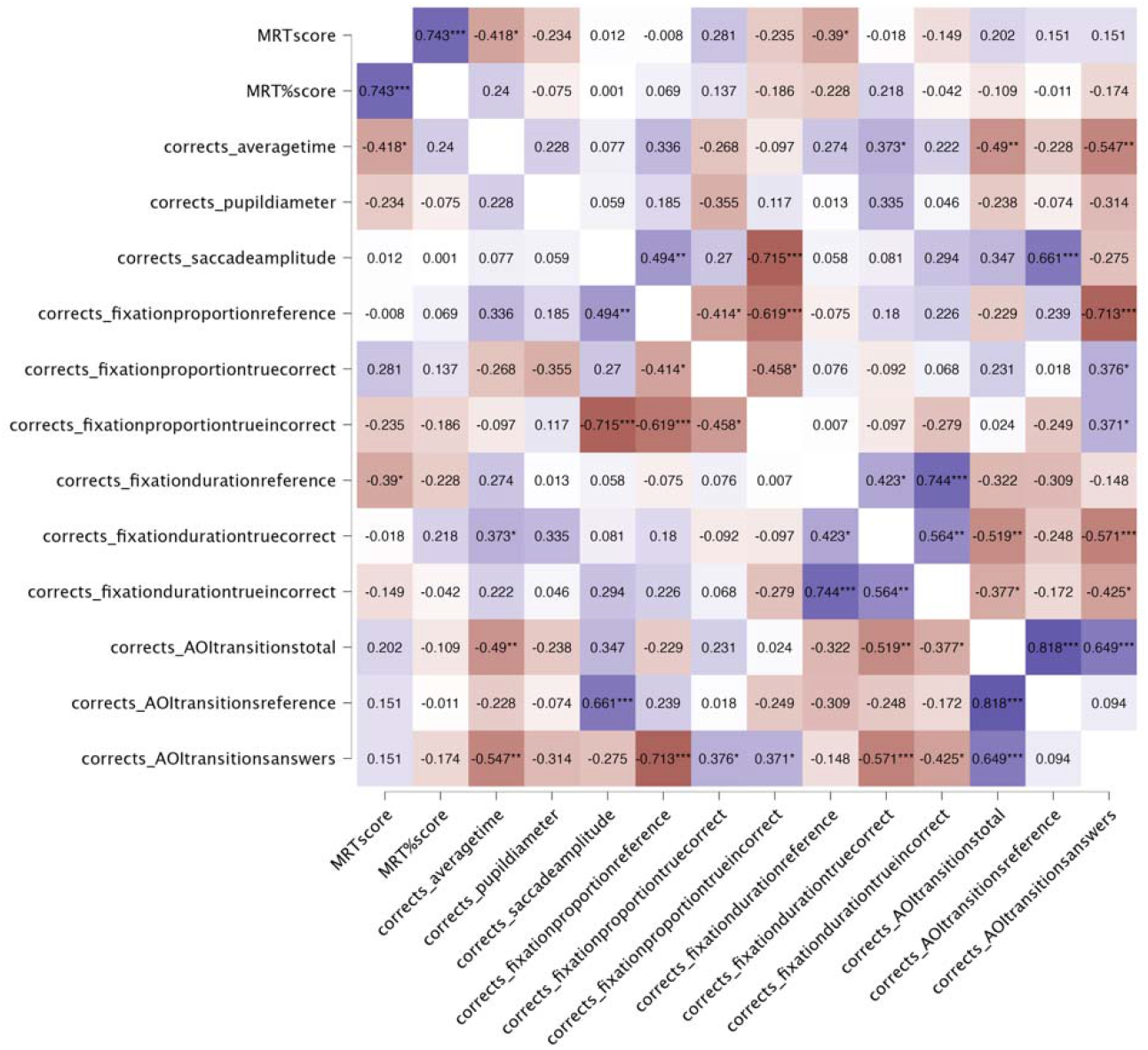
correlations with MRTscore, MRT%score, and all correct trial measures for all participants. Significant correlations are marked with asterisks (* p < .05, ** p < .01, *** p < .001). Positive correlations are shown in blue, negative correlations in red, with colour intensity proportional to magnitude of the coefficient.

In correct trials of all participants, *MRTscore* is negatively correlated with averagetime (*r* (28) = –.42, *p* = .022) and fixationdurationreference (*r* (28) = –.39, *p* = .033). No associations were found for *MRT%score*.

**Figure C2:**
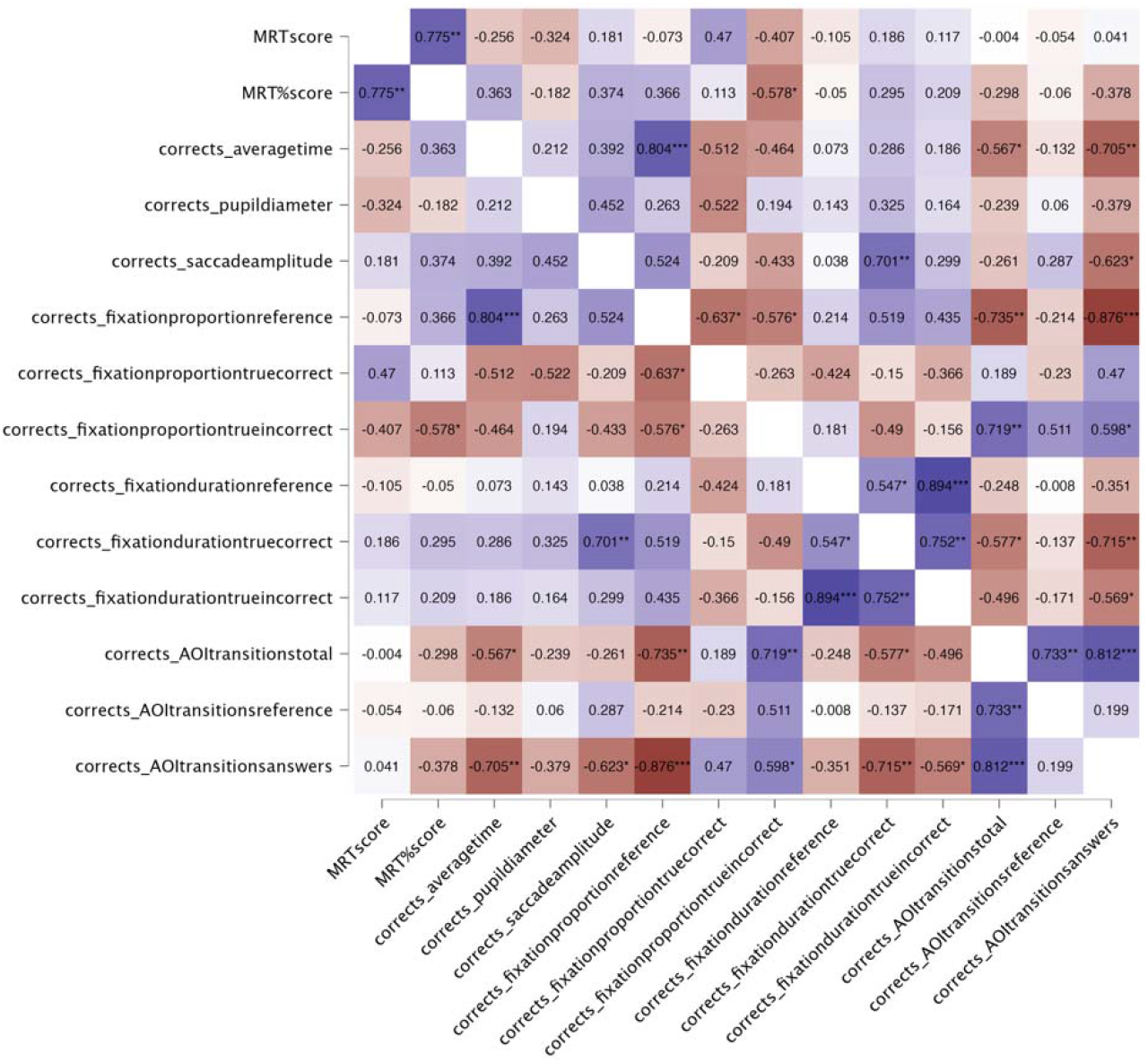
correlations with MRTscore, MRT%score, and all correct trial measures for female participants. Significant correlations are marked with asterisks (* p < .05, ** p < .01, *** p < .001). Positive correlations are shown in blue, negative correlations in red, with colour intensity proportional to magnitude of the coefficient.

In correct trials of female participants, *MRTscore* approaches positive correlation with *fixationproportiontruecorrect* (*r* (12) = .47, *p* = .090). MRT%score is negatively correlated with *fixationproportiontrueincorrect* (*r* (12) = –.58, *p* = .030).

**Figure C3:**
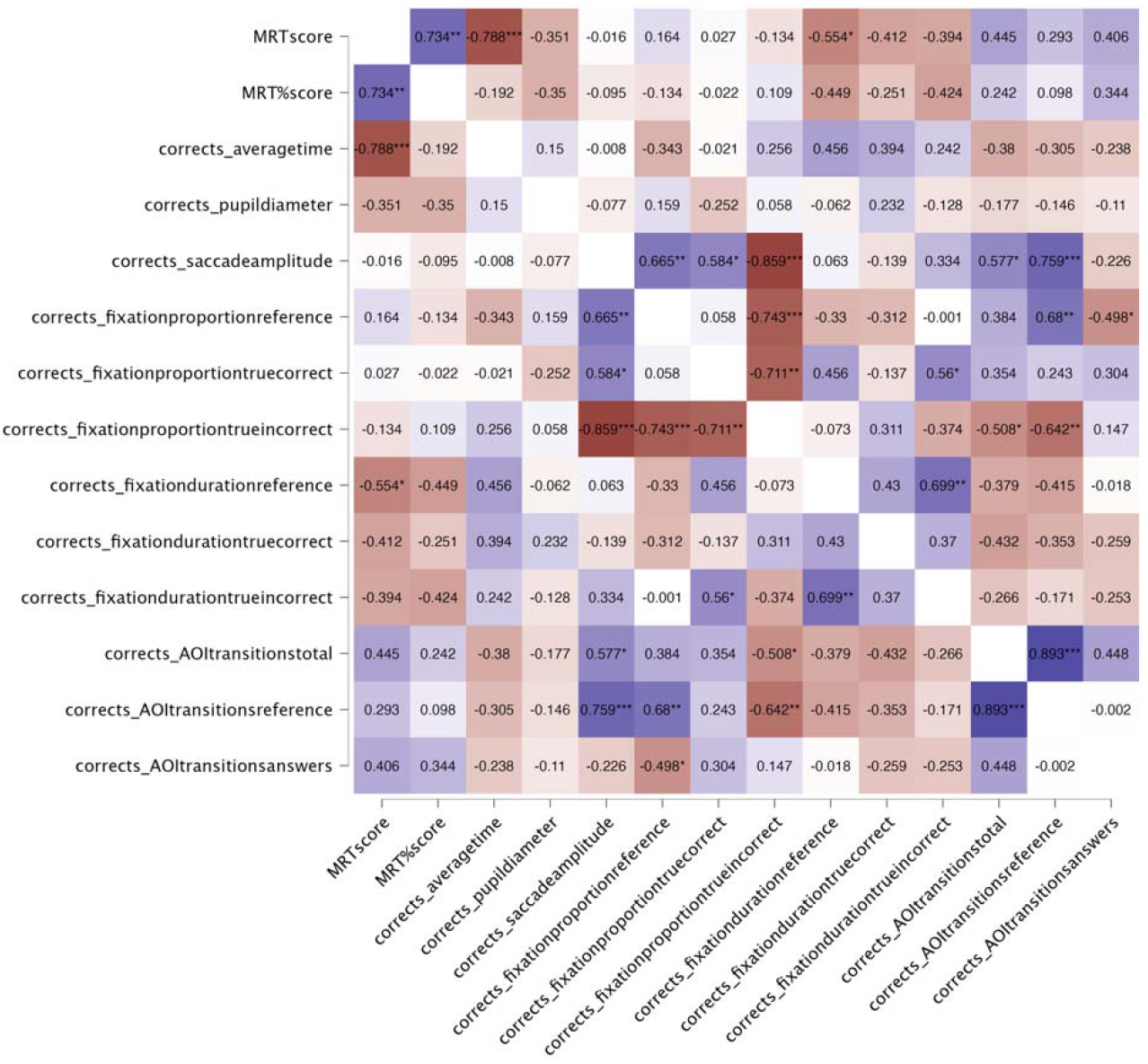
correlations with MRTscore, MRT%score, and all correct trial measures for male participants. Significant correlations are marked with asterisks (* p < .05, ** p < .01, *** p < .001). Positive correlations are shown in blue, negative correlations in red, with colour intensity proportional to magnitude of the coefficient.

In correct trials of male participants, *MRTscore* is negatively correlated with *averagetime* (*r* (14) = –.79, *p* < .001) and *fixationdurationreference* (*r* (14) = –.55, *p* = .026), and approaches positive correlation with *AOItransitionstotal* (*r* (14) = .45, *p* = .084). *MRT%score* approaches negative correlation with *fixationdurationreference* (*r* (14) = – .45, *p* = .081).

**Figure C4:**
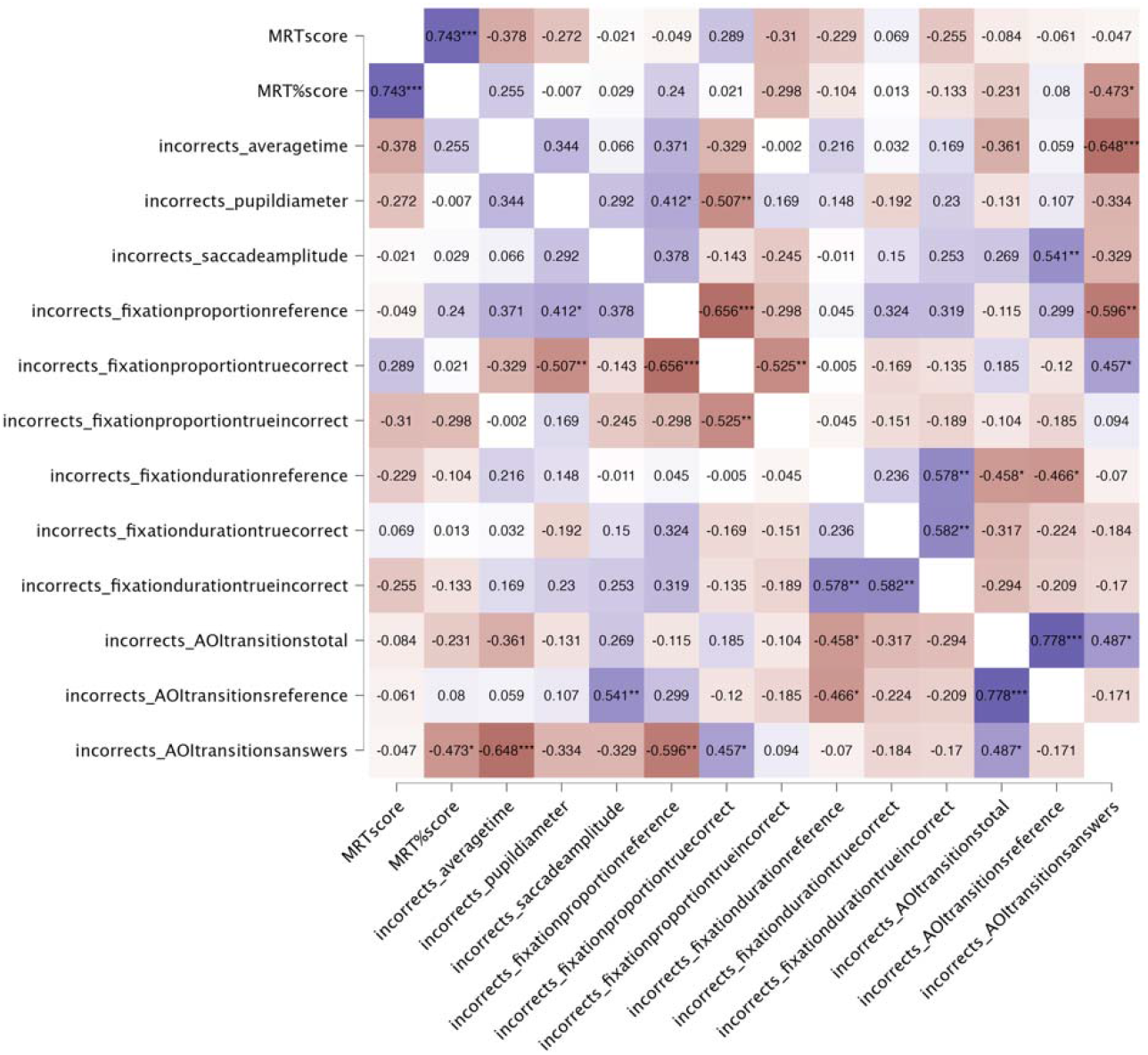
correlations with MRTscore, MRT%score, and all incorrect trial measures for all participants. Significant correlations are marked with asterisks (* p < .05, ** p < .01, *** p < .001). Positive correlations are shown in blue, negative correlations in red, with colour intensity proportional to magnitude of the coefficient.

In incorrect trials of all participants, *MRTscore* approaches negative correlation with *averagetime* (*r* (25) = –.38, *p* = .052). *MRT%score* is negatively correlated with *AOItransitionsanswers* (*r* (25) = –.47, *p* = .013).

**Figure C5:**
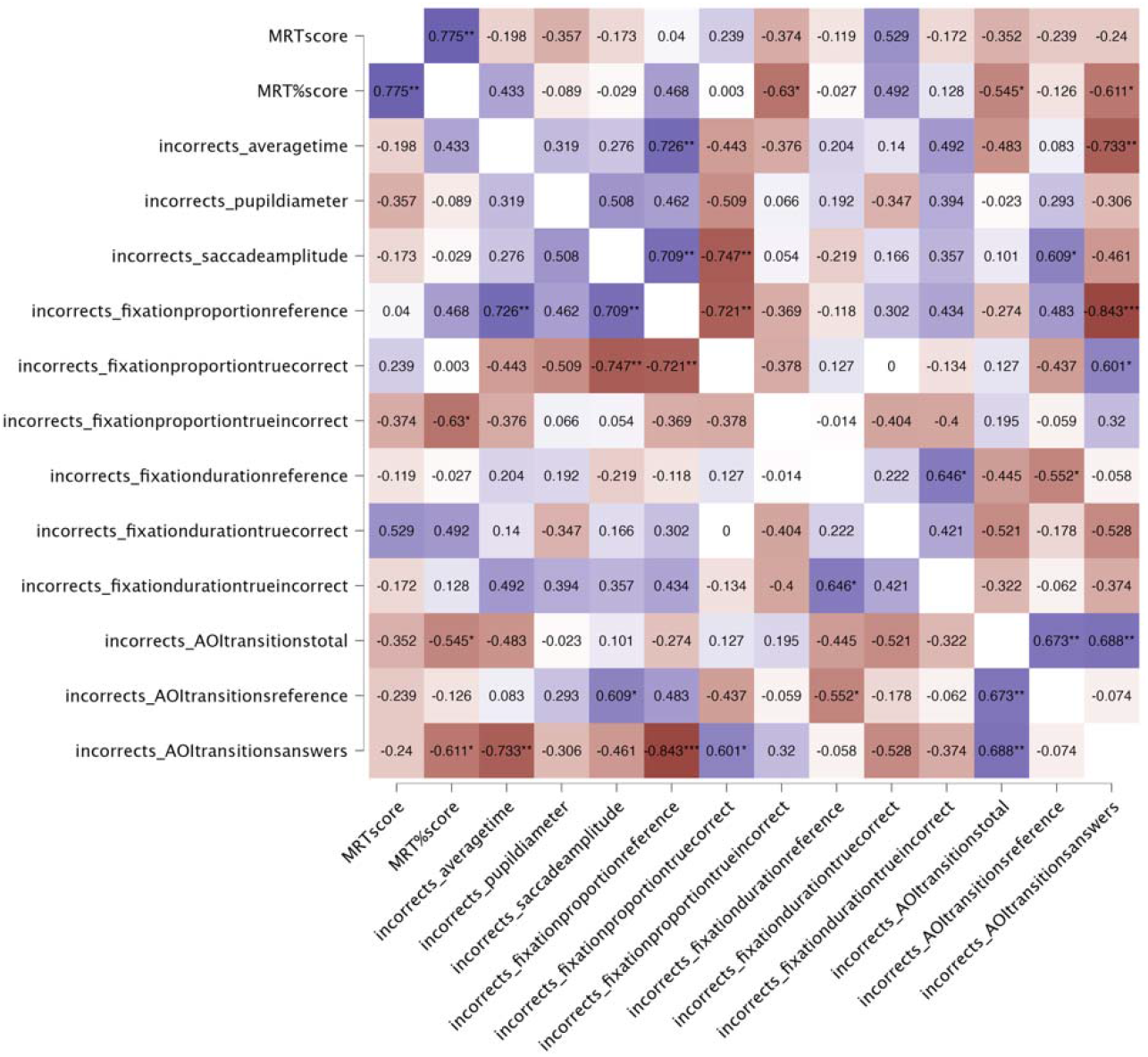
correlations with MRTscore, MRT%score, and all incorrect trial measures for female participants. Significant correlations are marked with asterisks (* p < .05, ** p < .01, *** p < .001). Positive correlations are shown in blue, negative correlations in red, with colour intensity proportional to magnitude of the coefficient.

In incorrect trials of female participants, *MRTscore* approaches positive correlation with *fixationdurationtruecorrect* (*r* (12) = .53, *p* = .052). *MRT%score* is negatively correlated with *fixationproportiontrueincorrect* (*r* (12) = –.63, *p* = .016), *AOItransitionstotal* (*r* (12) = –.55, *p* = .044), and *AOItransitionsanswers* (*r* (12) = –.61, *p* = .020); and approaches positive correlation with *fixationdurationtruecorrect* (*r* (12) = .49, *p* = .074) and *fixationproportionreference* (*r* (12) = .47, *p* = .091).

**Figure C6:**
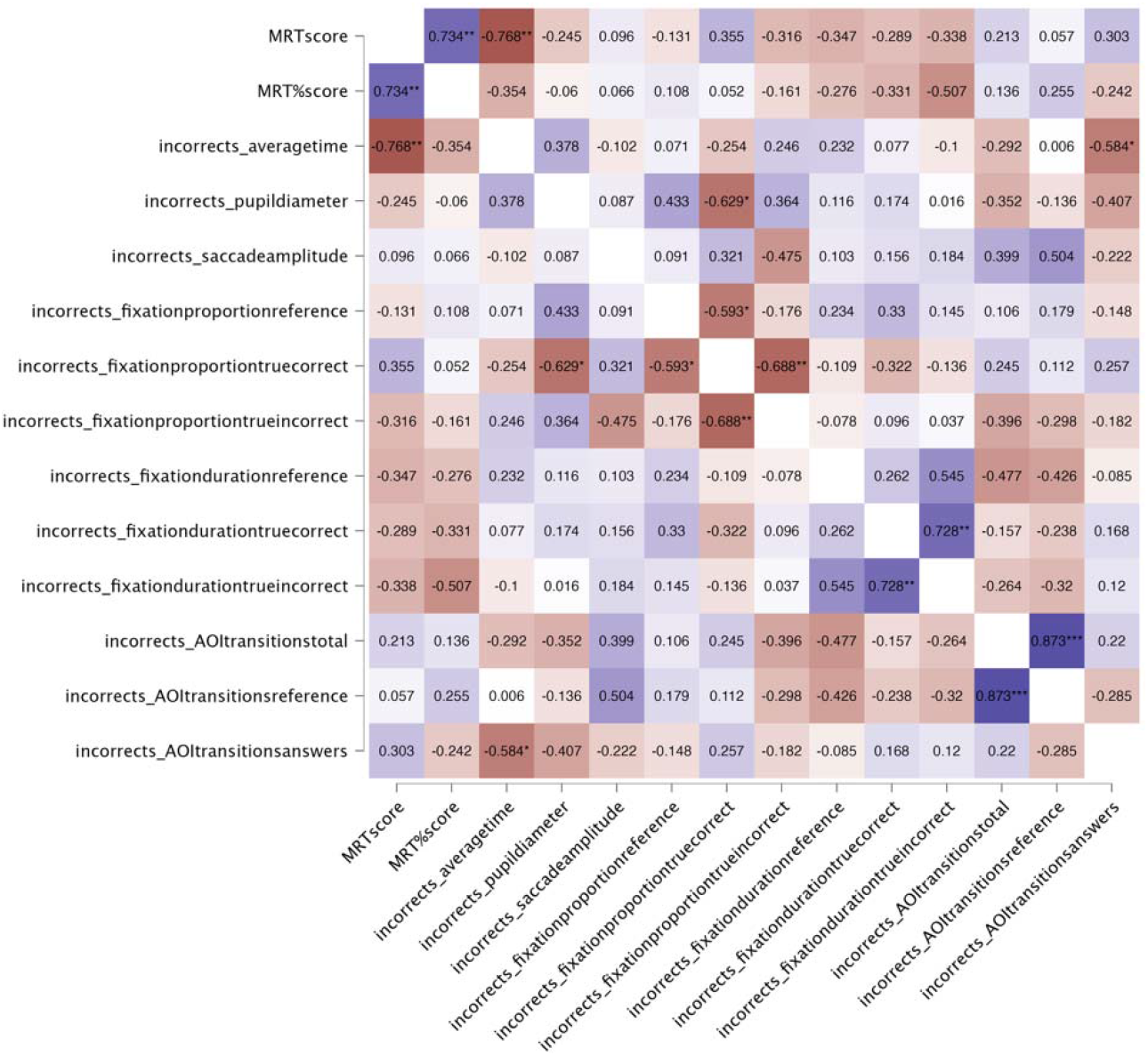
correlations with MRTscore, MRT%score, and all incorrect trial measures for male participants. Significant correlations are marked with asterisks (* p < .05, ** p < .01, *** p < .001). Positive correlations are shown in blue, negative correlations in red, with colour intensity proportional to magnitude of the coefficient.

In incorrect trials of male participants, *MRTscore* is negatively correlated with *averagetime* (*r* (11) = –.77, *p* = .002). *MRT%score* approaches negative correlation with *fixationdurationtrueincorrect* (*r* (11) = –.51, *p* = .077).

